# Molecular and structural mechanisms of ZZ domain-mediated cargo recognition by autophagy receptor Nbr1

**DOI:** 10.1101/2021.02.07.430097

**Authors:** Ying-Ying Wang, Jianxiu Zhang, Xiao-Man Liu, Meng-Qiu Dong, Keqiong Ye, Li-Lin Du

**Author notes:** These authors contributed equally (Ying-Ying Wang, Jianxiu Zhang). Lead contact: Li-Lin Du.

## Abstract

In selective autophagy, cargo selectivity is determined by autophagy receptors. However, it remains scarcely understood how autophagy receptors recognize specific protein cargos. In the fission yeast *Schizosaccharomyces pombe*, a selective autophagy pathway termed Nbr1-mediated vacuolar targeting (NVT) employs Nbr1, an autophagy receptor conserved across eukaryotes including humans, to target cytosolic hydrolases into the vacuole. Here, we identify two new NVT cargos, the mannosidase Ams1 and the aminopeptidase Ape4, that bind competitively to the first ZZ domain of Nbr1 (Nbr1-ZZ1). High-resolution cryo-EM analyses reveal how a single ZZ domain recognizes two distinct protein cargos. Nbr1-ZZ1 not only recognizes the N-termini of cargos via a conserved acidic pocket, similar to other characterized ZZ domains, but also engages additional parts of cargos in a cargo-specific manner. Our findings unveil a single-domain bispecific mechanism of autophagy cargo recognition, elucidate its underlying structural basis, and expand the understanding of ZZ domain-mediated protein-protein interactions.

## Introduction

Autophagy can transport cytoplasmic materials into lysosomes or vacuoles in selective manners. In recent years, a growing number of selective autophagy pathways have been uncovered (Stolz et al., 2014; Gatica et al., 2018). In these pathways, cargos are recognized by selectivity factors termed autophagy receptors. Autophagy receptors have been classified into three types based on their modes of cargo recognition (Behrends and Fulda, 2012; Zhao et al., 2020). Type I receptors bind cargo proteins directly via cargo-specific binding domains. Type II receptors recognize ubiquitin modifications on cargos via ubiquitin-binding domains. Type III receptors, which confer selectivity to organelle autophagy pathways, are integral membrane proteins embedded in the membranes of cargo organelles. Type I receptors, in particular, offer high levels of specificity and versatility in cargo recognition, but relatively little is known about the molecular and structural details of the interactions between type I receptors and their cargos (Kim et al., 2016; Yamasaki et al., 2016).

In the fission yeast *Schizosaccharomyces pombe,* we previously discovered a noncanonical selective autophagy pathway, Nbr1-mediated vacuolar targeting (NVT), in which the autophagy receptor Nbr1 cooperates with the endosomal sorting complexes required for transport (ESCRTs) to transport two aminopeptidases Lap2 and Ape2 from the cytosol into the vacuole (Liu et al., 2015). In mammalian cells, NBR1, a homolog of *S. pombe* Nbr1, promotes autophagic turnover of protein aggregates, peroxisomes, midbodies, and focal adhesions (Kirkin et al., 2009; Kuo et al., 2011; Deosaran et al., 2013; Kenific et al., 2016). In the budding yeast *Saccharomyces cerevisiae*, Atg19, a protein evolutionarily related to Nbr1 despite a lack of sequence homology (Kraft et al., 2010), is the autophagy receptor of a selective autophagy pathway termed the cytoplasm-to-vacuole targeting (Cvt) pathway (Scott et al., 2001). Cargos transported in an Atg19-dependent manner include three aminopeptidases Ape1, Ape4, and Lap3, an α-mannosidase Ams1, and virus-like particles of the Ty1 transposon (Kageyama et al., 2009; Suzuki et al., 2011; Shintani et al., 2002; Yuga et al., 2011). Both *S. pombe* Nbr1 and *S. cerevisiae* Atg19 are type I autophagy receptors that recognize cargos by direct protein-protein interactions (Liu et al., 2015; Yamasaki and Noda, 2017). They are ideal models to investigate how selective cargo recognition is achieved, and in particular, how a single receptor recognizes multiple cargos. However, thus far, only the Atg19-Ape1 interaction has been characterized at atomic resolution (Yamasaki et al., 2016; Yamasaki and Noda, 2017).

*S. pombe* Nbr1 has three ZZ domains (Liu et al., 2015; Kraft et al., 2010). Its homologs in diverse eukaryotic organisms, including human NBR1, also possess one or multiple ZZ domains (Kraft et al., 2010). ZZ domains are approximately 50-amino-acid long domains containing two structural zinc ions (Ponting et al., 1996; Legge et al., 2004; Zhang et al., 2019), and are known to mediate protein-protein interactions (Liu et al., 2015; Cha-Molstad et al., 2017; Kwon et al., 2018; Zhang et al., 2018a; Mi et al., 2018; Zhang et al., 2018b; Liu et al., 2020; Danielsen et al., 2012). Our previous study showed that the second and the third ZZ domains (ZZ2-ZZ3) of Nbr1 together mediate its interactions with NVT cargos Lap2 and Ape2 (Liu et al., 2015). Whether the first ZZ domain (ZZ1) of Nbr1 also functions in cargo recognition is unclear.

Recently, several atomic structures of ZZ domains in complex with ligand peptides have been solved by X-ray crystallography or NMR spectroscopy, including the ZZ domain of p62/SQSTM1 bound to N-degron peptides (Kwon et al., 2018; Zhang et al., 2018b), the ZZ domains of p300 and ZZZ3 bound to an N-terminal peptide of histone H3 (Zhang et al., 2018a; Mi et al., 2018), and the ZZ domain of HERC2 bound separately to N-terminal peptides of histone H3 and SUMO (Liu et al., 2020). These structures revealed a common theme of ZZ-mediated binding: ZZ domains bind the N-termini of their interacting proteins. Because these structural studies used only short peptides corresponding to the first few amino acids of ZZ-interacting proteins, it remains unclear whether and how ZZ domains bind other parts of their interactors.

Here, we identify *S. pombe* Ams1 and Ape4, orthologs of two known cargos of the Cvt pathway in *S. cerevisiae*, as additional cargos of the NVT pathway. The ZZ1 domain of Nbr1 bi-specifically binds these two cargo proteins and is required for their vacuolar targeting. We obtain atomic-resolution cryo-EM structures of the Ams1-ZZ1 complex and the Ape4-ZZ1 complex. These structures show that Nbr1-ZZ1 uses a conserved binding pocket to recognize the N-termini of Ams1 and Ape4 in a manner resembling the known mode of ZZ-mediated binding. Importantly, this binding mode is necessary but not sufficient. We find that Nbr1-ZZ1 also engages non-N-terminal regions of cargo proteins using cargo-specific binding surfaces, thus revealing unexpected sophistication and complexity of ZZ-mediated binding. Our study enriches the knowledge on how autophagy receptors achieve cargo selection and broadens the mechanistic understanding of ZZ domain-mediated interactions.

## Results

### Nbr1 mediates vacuolar targeting of Ams1 and Ape4

In an affinity purification coupled with mass spectrometry (AP-MS) analysis using *S. pombe* Nbr1 as bait, we found that, in addition to Lap2 and Ape2, two other soluble hydrolases co-purified with Nbr1—one is the α-mannosidase SPAC513.05/Ams1, the ortholog of *S. cerevisiae* Ams1; the other is the aminopeptidase SPAC4F10.02 (named Ppp20 in a previous publication (Idiris et al., 2006) and named Aap1 in PomBase), which we renamed Ape4 because it is the ortholog of *S. cerevisiae* Ape4 (Figure 1A). It is noteworthy that current database annotation of Ape4 misses its first six amino acids (Figure S1A). We confirmed both the Nbr1-Ams1 interaction and the Nbr1-Ape4 interaction using co-immunoprecipitation (Figure 1B and 1C).

**Figure 1.**
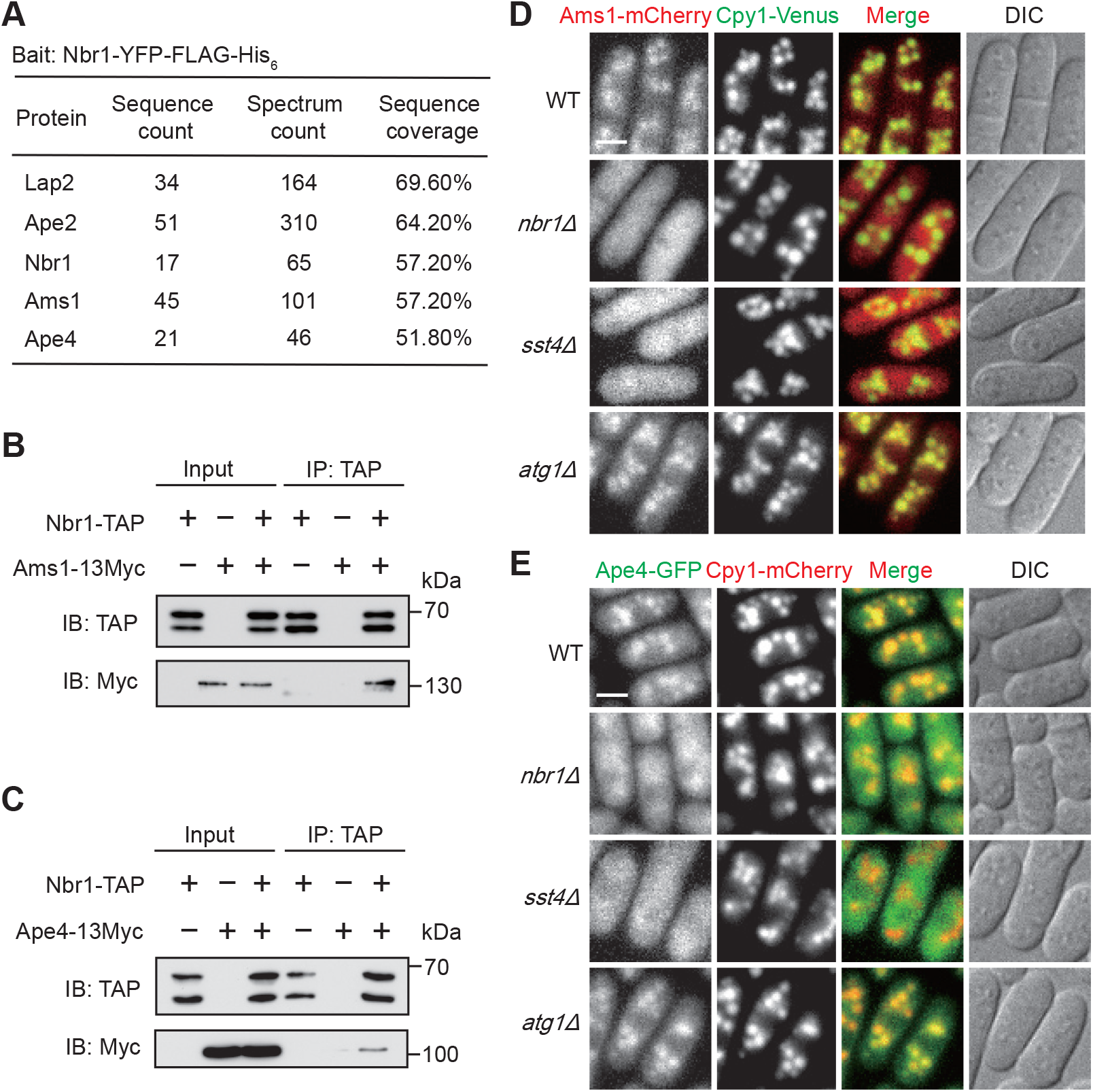
Ams1 and Ape4 are Nbr1 interactors and are cargos of the NVT pathway. (A) Identification of Ams1 and Ape4 as Nbr1 interactors. Nbr1-associated proteins were identified by affinity purification coupled with mass spectrometry (AP-MS) analysis using YFP-FLAG-His_6_-tagged Nbr1 as bait. (B-C) Coimmunoprecipitation of Ams1 (B) and Ape4 (C) with Nbr1. Endogenously TAP-tagged Nbr1 was immunoprecipitated (IP) using IgG Sepharose beads. Cell lysates and immunoprecipitates were examined by immunoblotting (IB). (D-E) Subcellular localization of Ams1 (D) and Ape4 (E). Wild-type (WT), *nbr1Δ*, *sst4Δ*, and *atg1Δ* cells were grown to mid-log phase in EMM medium and examined by live cell imaging. In (D), Ams1 was endogenously tagged at its C-terminus with mCherry. In (E), Ape4 was endogenously tagged at its C-terminus with GFP and in the same cells GFP-binding-protein (GBP) fused 3xUb was expressed under the *P41nmt1* promoter. Cpy1 is a vacuole lumen marker. Bar, 3 μm.

Like their budding yeast orthologs, fission yeast Ams1 and Ape4 do not possess signal peptides and thus are not expected to enter the secretory pathway. Nonetheless, C-terminally mCherry-tagged Ams1 exhibited a vacuole lumen localization (Figure 1D). This localization was abolished in *nbr1Δ* cells (Figure 1D). Moreover, immunoblotting analysis showed that Ams1 tagged with mECitrine (a GFP variant) was processed in a vacuolar protease-dependent manner and this processing was dependent on Nbr1 (Figure S1B). Together, these data indicate that Ams1 is targeted to the vacuole lumen by Nbr1.

For Ape4, we did not observe an obvious vacuolar localization above the cytosolic level (Figure S1C), possibly because its vacuolar targeting efficiency is lower than other NVT cargos. Given that ubiquitination promotes NVT transport (Liu et al., 2015), we adopted the method of artificial ubiquitin tethering to enhance the transport efficiency of individual cargos (Zhu et al., 2017). When three tandemly linked ubiquitin (3xUb) fused with the GFP-binding protein (GBP) was expressed in the cell, GFP-tagged Ape2, a known NVT cargo, localized more prominently to the vacuole, and Ub tethering-enhanced vacuolar targeting of Ape2 still required Nbr1 (Figure S1D and S1E). Applying this method to Ape4 resulted in an obvious vacuole lumen localization of Ape4 in wild-type but not in *nbr1Δ* cells (Figure 1E).

If Ams1 and Ape4 are cargos of the NVT pathway, the ESCRT machinery but not the macroautophagy machinery should be required for their vacuolar targeting. Indeed, Ams1 no longer localized in the vacuole in cells lacking *sst4*, which encodes a subunit of the ESCRT-0 complex (Figure 1D). In contrast, *atg1Δ* mutant showed a normal localization of Ams1 (Figure 1D). In addition, Ams1-mECitrine processing was blocked by *sst4Δ* but not *atg5Δ* (Figure S1F and S1G). Similar to the situation of Ams1, Ub tethering-enhanced Ape4 localization in the vacuole required Sst4 but not Atg1 (Figure 1E). These results demonstrate that Ams1 and Ape4 are NVT cargos.

### Vacuolar targeting of Ams1 and Ape4 is independent of Lap2 and Ape2

The previously identified NVT cargos, Lap2 and Ape2, require each other for interacting with Nbr1 and for vacuolar targeting (Liu et al., 2015). We wondered whether vacuolar targeting of Ams1 and Ape4 also depends on other cargos. The vacuole lumen localization of Ams1 and Ape4 was unaltered when Lap2 or Ape2 was absent (Figure 2A and 2B). In addition, Ams1 and Ape4 did not require each other for vacuolar targeting (Figure 2A and 2B). Furthermore, the absence of Ams1 or Ape4 had no effect on vacuolar targeting of Nbr1, Lap2, and Ape2 (Figure S2A-S2C). Thus, the mutual dependency between NVT cargos is limited to Lap2 and Ape2.

**Figure 2.**
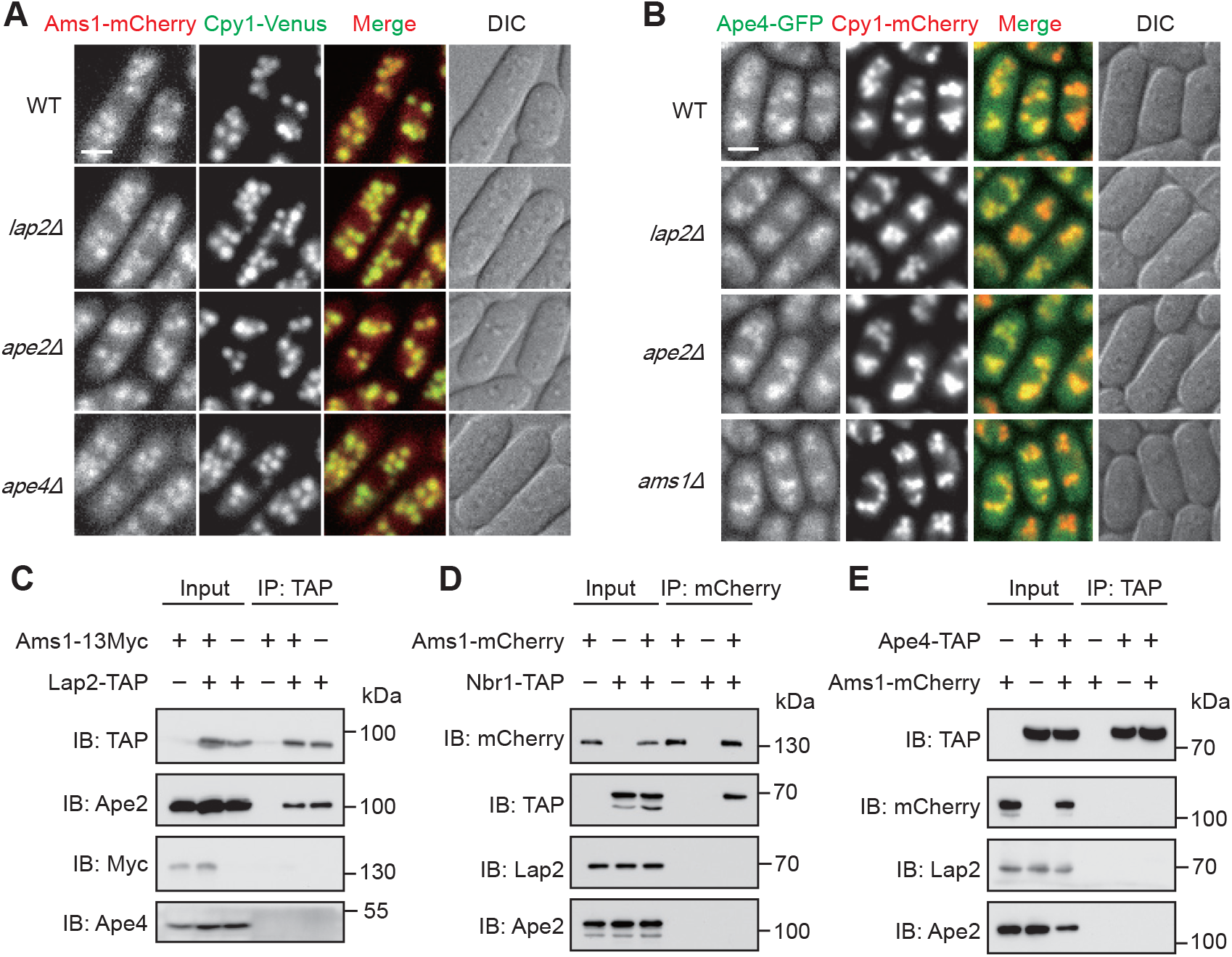
Ams1 and Ape4 do not rely on other cargos for vacuolar targeting and do not interact with other cargos. (A) Localization of Ams1 in WT, *lap2Δ*, *ape2Δ*, and *ape4Δ* cells. Bar, 3 μm. (B) Localization of Ape4 in WT, *lap2Δ*, *ape2Δ*, and *ams1Δ* cells. Bar, 3 μm. (C) Ape2 but not Ams1 or Ape4 co-immunoprecipitated with Lap2. Endogenously TAP-tagged Lap2 was immunoprecipitated. Ams1 was endogenously tagged with a 13Myc tag. (D) Nbr1 but not Lap2 or Ape2 co-immunoprecipitated with Ams1. Endogenously mCherry-tagged Ams1 was immunoprecipitated. Nbr1 was endogenously tagged with a TAP tag. (E) Ams1, Lap2, and Ape2 did not co-immunoprecipitate with Ape4. Endogenously TAP-tagged Ape4 was immunoprecipitated. Ams1 was endogenously tagged with an mCherry tag.

As we have previously shown that Nbr1, Lap2, and Ape2 associate together to form a complex (Liu et al., 2015), we investigated whether Ams1 and Ape4 are part of this complex. Ape2 but not Ams1 or Ape4 co-immunoprecipitated with Lap2 (Figure 2C). Consistent with this result, in AP-MS analyses, Ams1 and Ape4 did not co-purify with Lap2 or Ape2 (Figure S2D and S2E). In addition, Nbr1 but not Lap2 or Ape2 co-immunoprecipitated with Ams1 (Figure 2D). Furthermore, Ape4 showed no interaction with Lap2, Ape2, or Ams1 (Figure 2E). Together, these results indicate that NVT cargos form three separate complexes with Nbr1: the Lap2-Ape2-Nbr1 complex, the Ams1-Nbr1 complex, and the Ape4-Nbr1 complex.

### ZZ domains of Nbr1 mediate cargo recognition

Nbr1 contains three ZZ domains (sequentially named ZZ1, ZZ2, and ZZ3) and a FW domain (also known as NBR1 domain) (Figure 3A) (Liu et al., 2015). Previously we showed that ZZ2 and ZZ3 together are sufficient for binding Lap2 and Ape2 (Liu et al., 2015). To systematically map the cargo-interacting regions of Nbr1, we employed an imaging-based method, termed Pil1 co-tethering assay, to examine pairwise protein-protein interactions (Pan et al., 2020). In this assay, we fuse a bait protein to Pil1, which forms filamentary structures. If a prey protein can interact with the Pil1-fused bait protein, it will exhibit co-localization with the bait on filamentary structures. Analyses with a series of Nbr1 fragments showed that the ZZ1 domain was necessary and sufficient for binding both Ams1 and Ape4 (Figure 3A, 3B, and S3). This was confirmed by co-immunoprecipitation (Figure 3C and 3D). Ams1 and Ape4 could bind either competitively or non-competitively to Nbr1-ZZ1. To distinguish the two possibilities, GST-fused Nbr1-ZZ1 was incubated with a mixture of Ams1 and Ape4 and pulled down with glutathione beads (Figure 3E). The amount of Ams1 in the pull-down was gradually reduced with increasing concentrations of Ape4. Moreover, overexpression of Ape4 in vivo markedly diminished vacuolar targeting of Ams1 (Figure 3F). These data indicate that Ams1 and Ape4 bind competitively to Nbr1-ZZ1.

**Figure 3.**
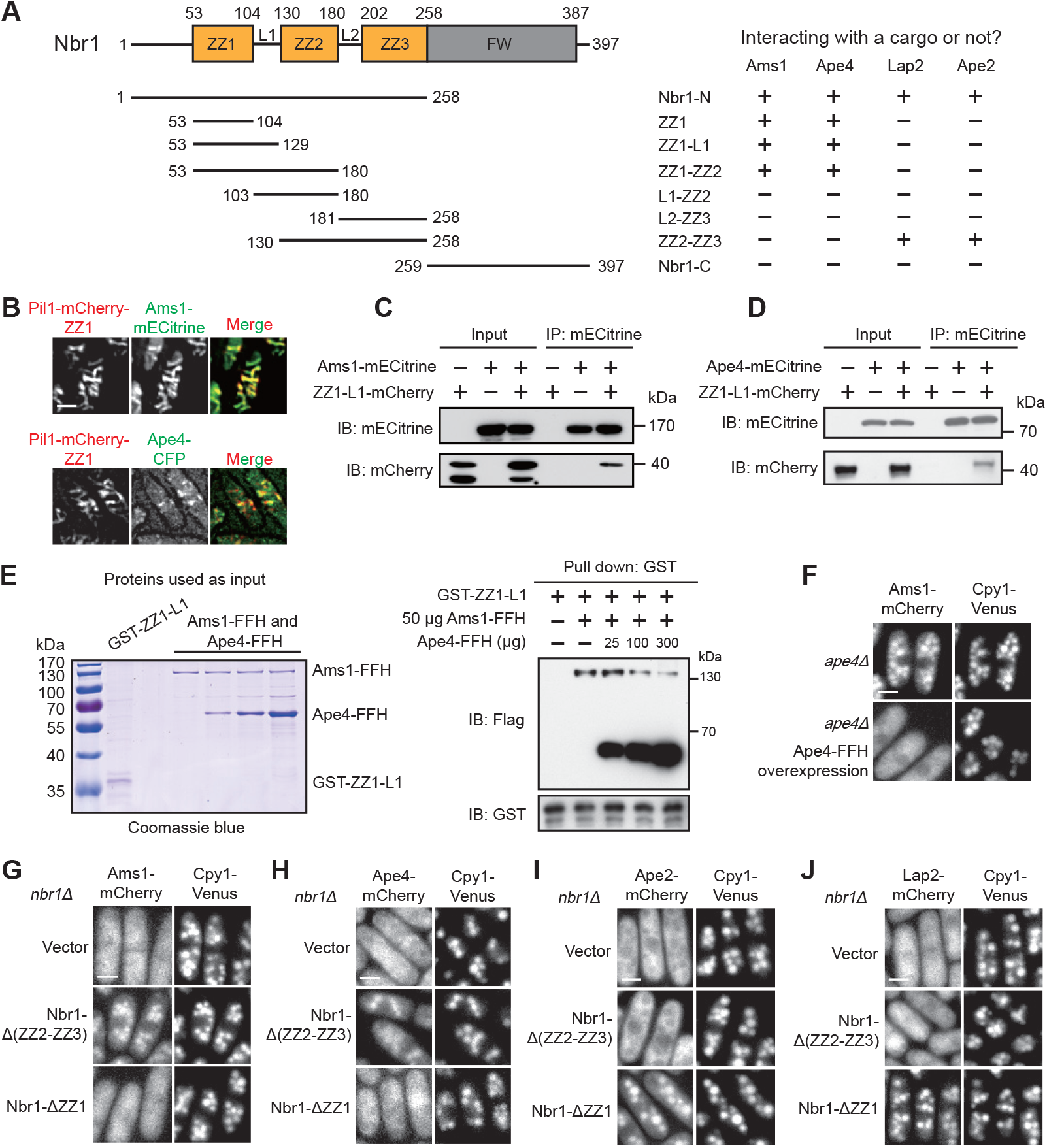
ZZ domains in Nbr1 are cargo recognition modules. (A) Truncation analysis revealed the regions of Nbr1 involved in its interactions with Ams1, Ape4, Lap2, and Ape2. Schematic at the top left shows the domain organization of Nbr1. Pil1 co-tethering assay data are shown in panel B and Supplementary Fig. 3. (B) The ZZ1 fragment of Nbr1 interacted with Ams1 and Ape4 in the Pil1 co-tethering assay. In this assay, a bait protein is fused to Pil1, which forms filamentary structures, and a prey protein that can interact with the bait exhibits co-localization on the same structures. Bar, 3 μm. (C) The ZZ1-L1 fragment of Nbr1 co-immunoprecipitated with Ams1. (D) The ZZ1-L1 fragment of Nbr1 co-immunoprecipitated with Ape4. (E) Ape4 competes with Ams1 for Nbr1 binding. GST-tagged ZZ1-L1 fragment of Nbr1 was bound to glutathione beads and incubated with FLAG_2_-His_6_ (FFH)-tagged Ams1 proteins and different amount of FFH-tagged Ape4. GST-tagged Nbr1 was purified from *E. coli*. FFH-tagged cargo proteins were purified from *S. pombe*. (F) Overexpressing of Ape4 inhibited vacuolar targeting of Ams1. FFH-tagged Ape4 was expressed under the strong *Pnmt1* promoter. Bar, 3 μm. (G) The ZZ1 region but not the ZZ2-ZZ3 region in Nbr1 is necessary for vacuolar targeting of Ams1. Bar, 3 μm. (H) The ZZ1 region but not the ZZ2-ZZ3 region in Nbr1 is necessary for vacuolar targeting of Ape4. Bar, 3 μm. (I) The ZZ2-ZZ3 region but not the ZZ1 region in Nbr1 is necessary for vacuolar targeting of Ape2. Bar, 3 μm. (J) The ZZ2-ZZ3 region but not the ZZ1 region in Nbr1 is necessary for vacuolar targeting of Lap2. Bar, 3 μm.

We next examined whether ZZ domain-mediated receptor-cargo interaction is important for vacuolar targeting of cargos. Nbr1 lacking ZZ2 and ZZ3, but not Nbr1 lacking ZZ1, rescued vacuolar targeting of Ams1 and Ape4 in *nbr1Δ* cells (Figure 3G and 3H). Conversely, Nbr1 lacking ZZ1, but not Nbr1 lacking ZZ2 and ZZ3, rescued vacuolar targeting of Lap2 and Ape2 in *nbr1Δ* cells (Figure 3I and 3J). Together with the interaction data, these results demonstrate that cargo recognition by Nbr1 is mediated by its ZZ domains, with one ZZ domain (ZZ1) bi-specifically recognizing two completely different cargos (the mannosidase Ams1 and the peptidase Ape4) and two other ZZ domains (ZZ2 and ZZ3) acting together to recognize the other two cargos Lap2 and Ape2.

### The cryo-EM structure of the Ams1-ZZ1 complex

To gain insights into how Nbr1 recognizes cargo proteins, we performed cryo-EM analyses on cargo-Nbr1 complexes. We first analyzed an Ams1-Nbr1 complex purified from *S. pombe*, but only resolved a structure of free Ams1 (Zhang et al., 2020). The interaction between Ams1 and Nbr1 appeared to be disrupted during cryo-EM sample preparation, possibly by exposure to the air-water interface (Glaeser and Han, 2017). To circumvent this problem, we fused a Nbr1 fragment encompassing ZZ1 and ZZ2 domains to the C-terminus of Ams1, with an intervening linker of 13 amino acids. The fusion protein was purified from *S. pombe* (Figure S4A) and imaged at a Titan Krios 300 kV microscope equipped with a K2 Summit camera. Following 2D and 3D classification, 227,295 particles were selected to reconstruct a density map at 2.4 Å overall resolution (Figure 4A and S4B-S4D). Ams1 and the ZZ1 domain of Nbr1 were resolved in the density map (Figure 4B and S5), whereas other portions of the fusion protein were invisible.

**Figure 4.**
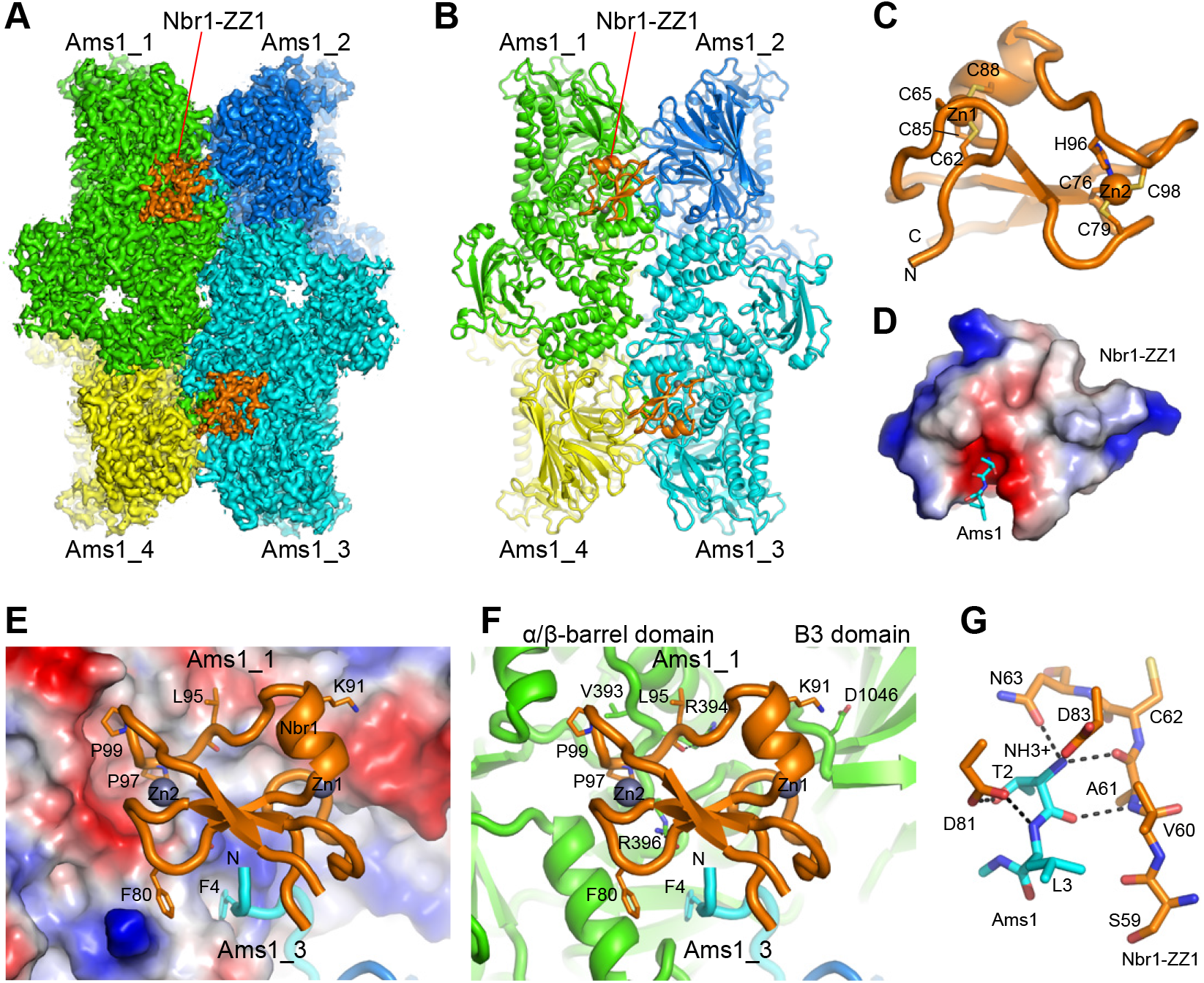
Cryo-EM structure of the Ams1-ZZ1 complex. (A) Cryo-EM density map of tetrameric Ams1 in complex with Nbr1-ZZ1 at 2.4 Å. Ams1 subunits 1 to 4 are colored in green, blue, cyan, and yellow. Nbr1-ZZ1 molecules are all colored in orange. (B) Ribbon representation of the Ams1-ZZ1 complex structure. (C) Structure of the Nbr1-ZZ1 domain. The N and C-termini and the Zn-coordinating residues are labeled. (D) Nbr1-ZZ1 uses an acidic pocket to bind the N-terminus of Ams1. Nbr1-ZZ1 is shown as an electrostatic potential surface where the positively to negatively charged regions are colored from blue to red. The N-terminal dipeptide of Ams1 are shown as sticks. The structure has the same orientation as C. (E-F) Binding interface between Nbr1-ZZ1 and Ams1. Nbr1-ZZ1 corresponds to the top left Nbr1-ZZ1 molecule in (A). Ams1_1 is shown as electrostatic potential surface (E) or ribbons (F). Ams1_3 (cyan) and Nbr1-ZZ1 (orange) are shown as ribbons. Residues involved in interactions are labeled. (G) Interactions between Nbr1-ZZ1 (orange) and the N-terminal dipepetide of Ams1 (cyan). Hydrogen bonds are denoted by black dashed lines.

Ams1 forms a tetramer in the complex structure (Figure 4A and 4B), as in the free Ams1 structure (Zhang et al., 2020). Nbr1-ZZ1 adopts a cross-brace zinc finger fold that coordinates two zinc ions and contains an N-terminal hairpin loop (a two-stranded β-sheet in some ZZ domains), a three-stranded antiparallel β-sheet, and an α-helix (Figure 4C), similar to other ZZ domain structures (Ponting et al., 1996; Cha-Molstad et al., 2017; Kwon et al., 2018; Zhang et al., 2018a; Mi et al., 2018; Zhang et al., 2018b; Liu et al., 2020). Four Nbr1-ZZ1 molecules bind symmetrically to one Ams1 tetramer (Figure 4A and 4B). Remarkably, each Nbr1-ZZ1 molecule simultaneously engages two Ams1 subunits through a continuous binding interface. Specifically, the Nbr1-ZZ1 molecule shown at the top left of Figure 4A and Figure 4B binds both the N-terminus of the Ams1_3 subunit (sub-interface I) and a large surface of the Ams1_1 subunit (sub-interface II) (Figure 4D-4F). This special binding configuration corresponds to the fact that the N-terminal tail of each Ams1 subunit extends away from the rest of the molecule and inserts into a cleft formed between two other Ams1 subunits (Zhang et al., 2020). Thus, the Ams1-ZZ1 complex represents an interesting example of a protein-binding module—in this case a ZZ domain—recognizing the quaternary structure of its oligomeric binding partner.

At sub-interface I, the N-terminal dipeptide (T2-L3) of Ams1 fits into an acidic pocket of Nbr1-ZZ1 located between the hairpin loop and the β-sheet (Figure 4D). The density map unambiguously shows that T2 is the most N-terminal residue of Ams1 (Figure S5E), consistent with the expectation that the first translated methionine residue is removed in vivo when the second residue has a small side chain (Bonissone et al., 2013). The N-terminal dipeptide of Ams1 aligns in an antiparallel manner with residues 60-62 of Nbr1 (Figure 4G). The free α-amino group of T2 hydrogen bonds with the side chain amide group of N63 and the carbonyl oxygen of A61, and forms an electrostatic interaction with the carboxylate group of D83 (Figure 4G). The carbonyl oxygen of T2 hydrogen bonds with the amide nitrogen of A61 and the amide nitrogen of L3 interacts with the side chain carboxylate group of D81 (Figure 4G). These interactions are directed to the backbone atoms of the dipeptide and hence are independent of peptide sequence. In addition, the hydroxyl group of T2 forms a hydrogen bond with the carboxylate group of D81 and the side chain of L3 makes van der Waals interactions with V60 of Nbr1 (Figure 4G). These interactions engage the side chains of the cargo and may confer sequence specificity.

At sub-interface II, Nbr1-ZZ1 binds the α/β barrel and the B3 domain of Ams1 (Figure 4E and 4F). Sub-interface II is much more extensive than sub-interface I. They bury solvent accessible surface areas of 760 Å^2^ and 227 Å^2^ per molecule, respectively. The binding at sub-interface II is mediated by shape complementarity and a large number of van der Waals, hydrophobic, hydrogen bonding, and electrostatic interactions (Figure 4E and 4F). In particular, the loop between P97 and P99 of Nbr1 fits snugly into a cavity on the surface of Ams1 (Figure 4E).

The entire structure of Ams1, including the N-terminal dipeptide, does not change between the free and ZZ1-bound states (Figure S5F), indicating that Nbr1-ZZ1 recognizes a preformed conformation of Ams1.

### The cryo-EM structure of the Ape4-ZZ1 complex

We applied the same fusion strategy to determine the structure of Ape4 in complex with Nbr1-ZZ1. The fusion of Ape4 and Nbr1(53-129) was expressed and purified from *S. pombe*, and analyzed by cryo-EM (Figure 5A and S6). A density map was reconstructed from 352,316 particles at 2.26 Å resolution (Figure S7).

**Figure 5.**
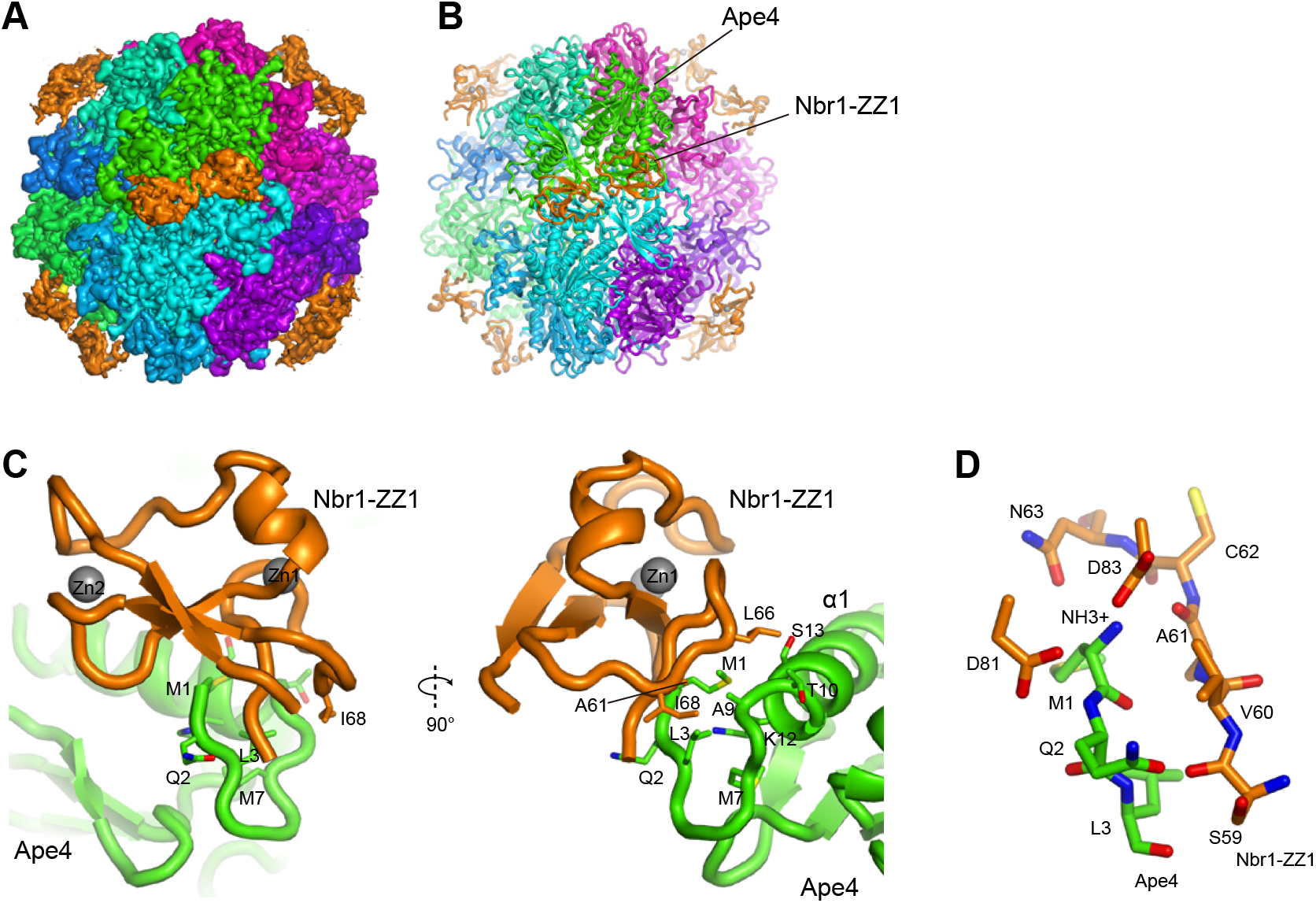
Cryo-EM structure of the Ape4-ZZ1 complex. (A) LocScale cryo-EM density map of Ape4 dodecamer in complex with Nbr1-ZZ1. Twelve Ape4 subunits are colored differently and all Nbr1-ZZ1 molecules are colored in orange. (B) Ribbon representation of the Ape4-ZZ1 complex structure. (C) Binding interface between Nbr1-ZZ1 and Ape4 shown in two orthogonal views. Residues involved in interactions are labeled. The ZZ1 domain in the left panel has the same orientation as that in Figure 4F. (D) Interactions between Nbr1-ZZ1 and the N-terminal tripeptide of Ape4.

Like other M18 family metallopeptidases with solved structures, including the Cvt pathway cargo *S. cerevisiae* Ape1 (ScApe1) (Chen et al., 2012; Chaikuad et al., 2012; Sivaraman et al., 2012; Su et al., 2015; Yamasaki et al., 2016; Bertipaglia et al., 2016; Yamasaki and Noda, 2017), *S. pombe* Ape4 forms a tetrahedron-shaped homo-dodecamer, with each subunit composed of a proteolytic domain and a wing domain (Figure 5A, 5B, and S8A-8C). In the Ape4-ZZ1 complex structure, 12 Nbr1-ZZ1 molecules are arranged in six pairs that each bind around the dyad axes of the Ape4 dodecamer (Figure 5A and 5B). Nbr1-ZZ1 showed poorly resolved density at ~ 4 Å resolution (Figure S7A), in part because of its partial occupancy (Figure S6E). Two Nbr1-ZZ1 molecules in a pair are in contact with each other, raising the possibility that they bind Ape4 in a cooperative manner. However, 3D variability analysis revealed no correlation between the densities of the two Nbr1-ZZ1 molecules, suggesting that they rather bind independently (Figure S6E).

Unlike the situation of the Ams1-ZZ1 complex, in the Ape4-ZZ1 complex, each Nbr1-ZZ1 molecule only interacts with one Ape4 subunit. Nbr1-ZZ1 contacts the N-terminal U-shaped tail and the α1 helix of Ape4, burying a solvent accessible surface area of 359 Å^2^ per molecule (Figure 5C). The interface is rather small, accounting for the low occupancy of Nbr1 in the complex. The interface can be divided into two sub-interfaces. Sub-interface I largely corresponds to sub-interface I in the Ams1-ZZ1 complex, with the N-terminal tripeptide M1-Q2-L3 of Ape4 fitting into the acidic pocket of Nbr1-ZZ1 (Figure 5D). The N-terminal methionine of Ape4 is retained (Figure S7E), because N-terminal methionine excision does not happen when the second residue is glutamine (Bonissone et al., 2013). The fact that Nbr1-ZZ1 uses the same acidic pocket to bind the N-termini of Ams1 and Ape4 explains why these two cargos exhibit mutually exclusive binding to Nbr1-ZZ1. The sub-interface II is formed by the contact between the hairpin loop of Nbr1-ZZ1 and the α1 helix of Ape4, mainly via van der Waals interactions (Figure 5C).

In Ape4, the N-terminal U-shaped tail packs against the α1 helix by forming a hydrophobic cluster composed of M1, L3, M7 and the aliphatic part of K12 (Figure 5C). It is likely that these intra-molecular contacts help stabilize the conformation of the N-terminal tail and facilitate its interaction with Nbr1-ZZ1. The conformation of the N-terminal tail of Ape4 is not conserved in the structure of ScApe1 (Figure S8B) (Yamasaki et al., 2016), but judging from sequence conservation (Figure S1A), is likely conserved in Ape4 orthologs in other fission yeast species.

### Mutational analysis of Nbr1-ZZ1

The structures of ZZ1-cargo complexes showed that the side chains of N63 and D81 of Nbr1 (part of the acidic pocket) are involved in the recognition of the N-termini of Ams1 and Ape4. Both N63R and D81R mutations individually abolished the ability of Nbr1 to bind Ams1 and Ape4 (Figure 6A and 6C), and also abolished the ability of Nbr1 to mediate vacuolar targeting of Ams1 and Ape4 (Figure 6B and 6D). Thus, the N-terminal binding mode is important for cargo recognition by Nbr1-ZZ1.

**Figure 6.**
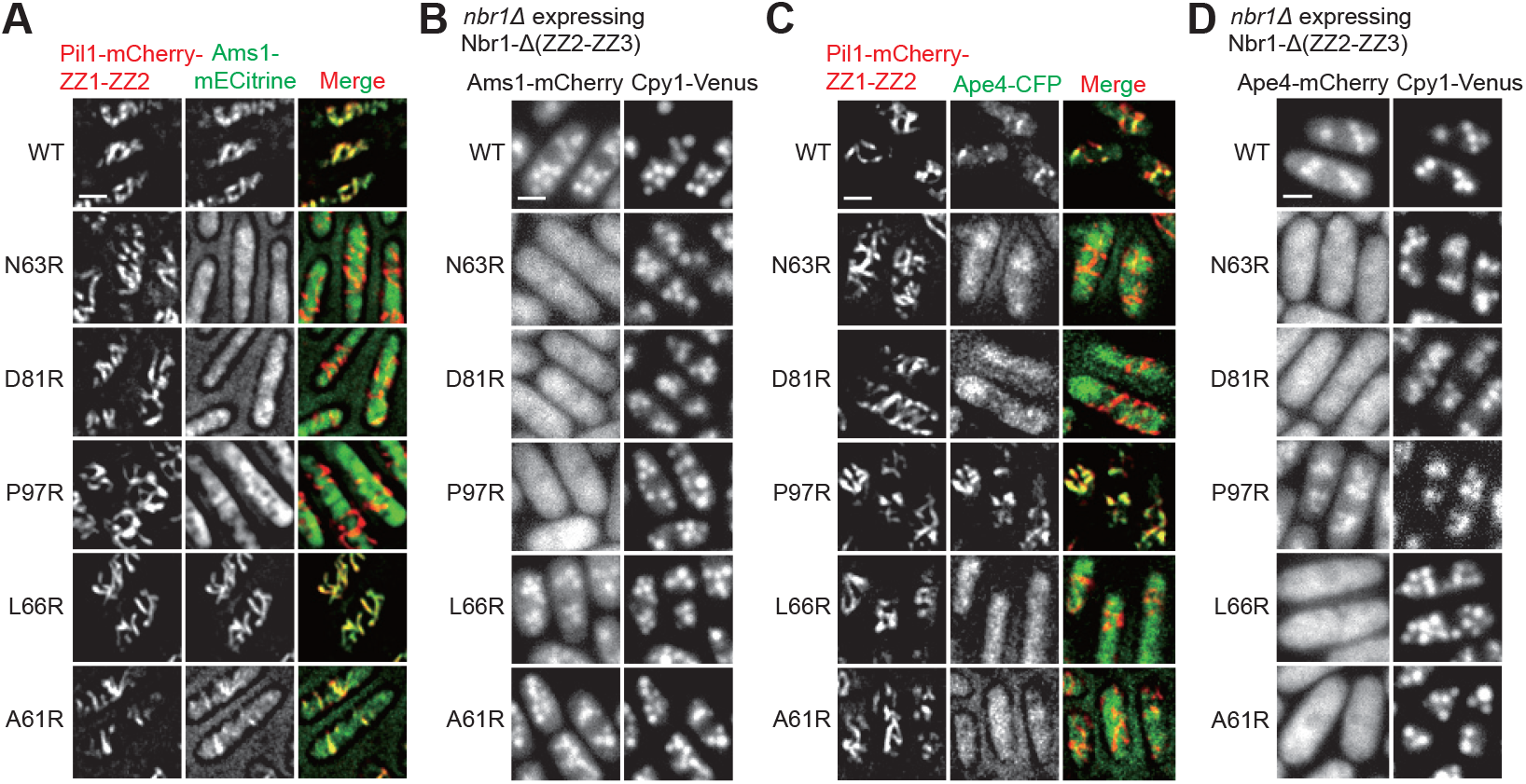
Mutational analysis of Nbr1-ZZ1 residues involved in cargo recognition. (A) The effects of Nbr1-ZZ1 mutations on its Ams1-binding ability. Nbr1-Ams1 interactions were examined by the Pil1 co-tethering assay. Mutations were introduced into the bait protein Pil1-mCherry-ZZ1-ZZ2. Bar, 3 μm. (B) The effects of Nbr1-ZZ1 mutations on its ability to mediate the vacuolar targeting of Ams1. Mutations were introduced into Nbr1-Δ(ZZ2-ZZ3), which was expressed in the *nbr1Δ* mutant background. Bar, 3 μm. (C) The effects of Nbr1-ZZ1 mutations on its Ape4-binding ability. Nbr1-Ape4 interactions were examined by the Pil1 co-tethering assay. Mutations were introduced into the bait protein Pil1-mCherry-ZZ1-ZZ2. Bar, 3 μm. (D) The effects of Nbr1-ZZ1 mutations on its ability to mediate the vacuolar targeting of Ape4. Mutations were introduced into Nbr1-Δ(ZZ2-ZZ3), which was expressed in the *nbr1Δ* mutant background. Bar, 3 μm.

Given that in the structures, Nbr1-ZZ1 uses cargo-specific binding surfaces to interact with non-N-terminal parts of cargos, we predicted that an Nbr1 mutation that disrupts a binding surface specific for one cargo may compromise the binding of that cargo but not the other cargo. Indeed, Nbr1-P97, which is situated at the Ams1-specific binding surface, when mutated to arginine, disrupted the Nbr1-Ams1 interaction (Figure 6A), and abolished the ability of Nbr1 to mediate vacuolar targeting of Ams1 (Figure 6B), but had no effect on the Nbr1-Ape4 interaction and the ability of Nbr1 to mediate vacuolar targeting of Ape4 (Figure 6C and 6D). Conversely, when we introduced into Nbr1 the L66R mutation, which affects a key residue at the Ape4-specific binding surface, the Nbr1-Ape4 interaction but not the Nbr1-Ams1 interaction was disrupted (Figure 6A and 6C), and vacuolar targeting of Ape4 but not Ams1 was abrogated (Figure 6B and 6D). The Nbr1-A61R mutation caused effects similar to those of the Nbr1-L66R mutation (Figure 6). Based on the structures, this mutation is expected to cause a steric clash with Ape4 but not with Ams1 (Figure 5C). These results demonstrate that cargo recognition by Nbr1-ZZ1 requires both its interactions with the N-termini of cargos and its interactions with non-N-terminal parts of cargos.

### Mutational analysis of Ams1 and Ape4

To verify the roles of the N-terminal residues of Ams1 and Ape4 in their recognition by Nbr1-ZZ1, we generated point mutations of the first two residues of Ams1 and the first three residues of Ape4. T2A and L3A mutations in Ams1 each diminished the Ams1-Nbr1 interaction (Figure 7A). For Ape4, to circumvent the effect of the M1A mutation in preventing translation and the effect of the Q2A mutation in causing the removal of M1, we fused the mature SUMO protein, Pmt3(1-111), to the N-terminus of Ape4, so that M1A and Q2A mutant forms of Ape4 can be generated through the cleavage of the fusion protein by the endogenous SUMO maturation enzyme Ulp1. As a control, we also fused Pmt3(1-109), which cannot be processed by Ulp1, to Ape4. Pmt3(1-111)-Ape4 but not Pmt3(1-109)-Ape4 exhibited an interaction with Nbr1 (Figure 7B). M1A, Q2A, and L3A mutations in Ape4 individually disrupted the Ape4-Nbr1 interaction (Figure 7B and 7C). As the side chains of M1 and L3 of Ape4 are sandwiched between the hairpin loop of Nbr1 and the α1 helix of Ape4 (Figure 5C), the sequence specificity for these two residues is probably determined by interactions from both Nbr1 and Ape4. The side chain of Q2 was not well resolved in the density map (Figure S7E), and it may be specified by D81 of Nbr1. Using the Pmt3(1-111) fusion approach, we also tested whether preserving M1 in Ams1 affects Nbr1 binding and found that N-terminal methionine removal is important for the Ams1-Nbr1 interaction (Figure 7D). Together, these results showed that the sequences of the N-terminal residues in both Ams1 and Ape4 are important for Nbr1-ZZ1 binding.

**Figure 7.**
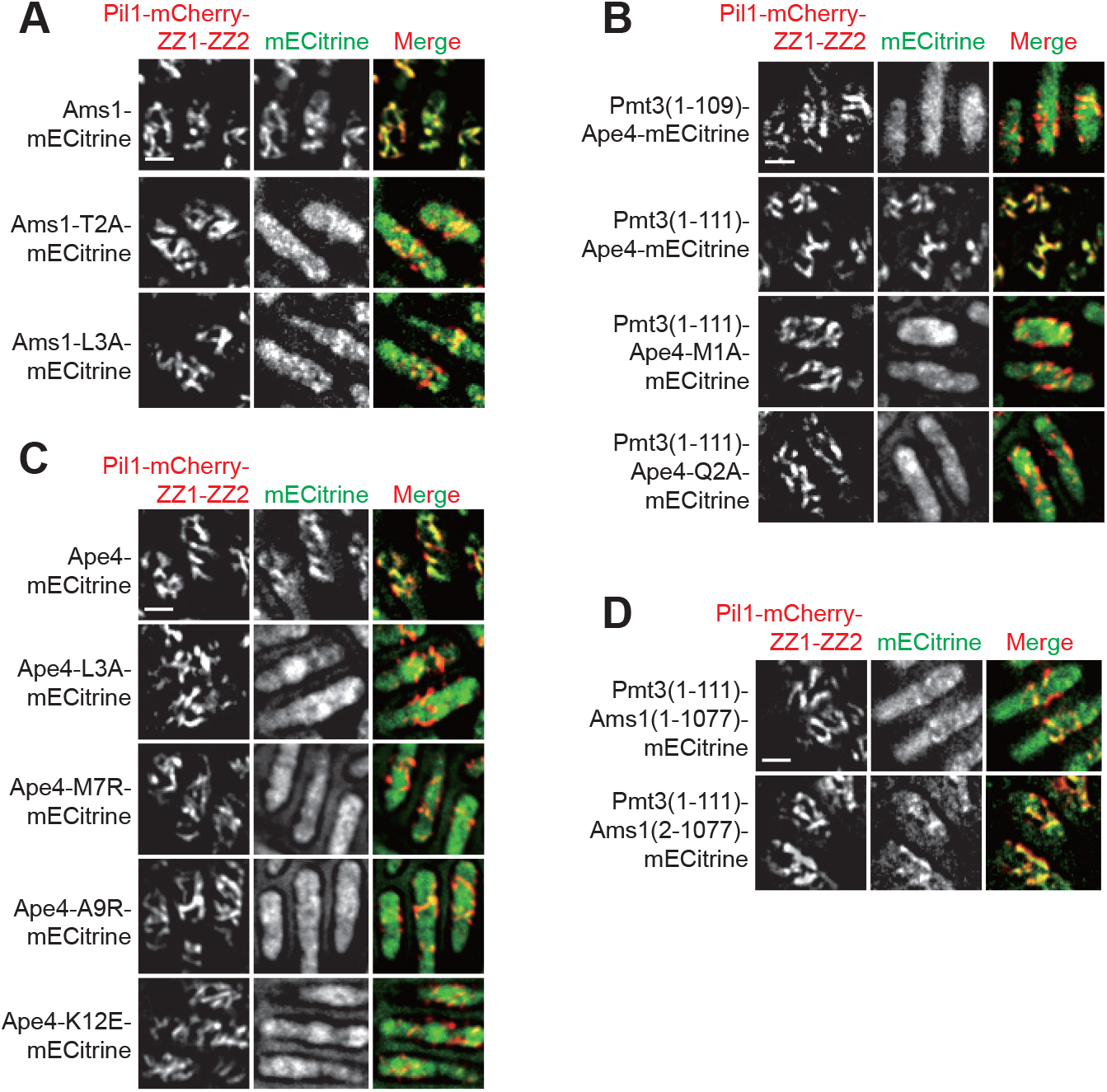
Mutational analysis of cargo residues involved in Nbr1 binding. (A) The effects of individually mutating the first two residues of Ams1 on its Nbr1-binding ability. Nbr1-Ams1 interactions were examined by the Pil1 co-tethering assay. Bar, 3 μm. (B) The effects of individually mutating the first two residues of Ape4 on its Nbr1-binding ability. Nbr1-Ape4 interactions were examined by the Pil1 co-tethering assay. Pmt3-Ape4 fusions were used to allow alanine substitutions to be introduced at M1 and Q2 positions. Bar, 3 μm. (C) The effects of individually mutating the third residue and other residues in the U-shaped loop of Ape4 on its Nbr1-binding ability. Nbr1-Ape4 interactions were examined by the Pil1 co-tethering assay. Bar, 3 μm. (D) The effect of preserving the M1 residue of Ams1 on its Nbr1-binding ability. Bar, 3μm.

To verify the role of the non-N-terminal binding mode in the Ape4-ZZ1 interaction, we introduced into Ape4 the A9R mutation, which was expected to cause a steric clash in the sub-interface II between Ape4 and Nbr1-ZZ1 (Figure 5C). Indeed, this mutation abolished the Ape4-Nbr1 interaction (Figure 7C). In addition, we found that the Ape4-M7R mutation and the Ape4-K12E mutation, which are expected to destabilize the conformation of the U-shaped tail of Ape4, also individually abolished the Ape4-Nbr1 interaction (Figure 7C). These data suggest that packing of the N-terminal tail to the structural body of Ape4 is important for Nbr1 binding.

## Discussion

*S. pombe* Nbr1 and *S. cerevisiae* Atg19, the autophagy receptors employed by the NVT pathway and the Cvt pathway respectively, do not share obvious sequence homology, but nonetheless have been shown by an evolutionary analysis to belong to the same protein family ubiquitously present in eukaryotes (Kraft et al., 2010). In this study, we identify Ams1 and Ape4 in *S. pombe* as cargos of the NVT pathway. Because their orthologs in *S. cerevisiae* are cargos of the Cvt pathway, our finding reinforces the functional similarity and the evolutionary relatedness of these two selective autophagy pathways that transport soluble hydrolases into the vacuole. Given that *S. pombe* and *S. cerevisiae* are two distantly related fungal species that diverged about 600 million years ago (Kumar et al., 2017), it is likely that Nbr1/Atg19-related proteins in other species, at least among fungi, are also autophagy receptors for hydrolase cargos.

Both *S. pombe* Nbr1 and *S. cerevisiae* Atg19 can recognize multiple hydrolase cargos. In *S. pombe* Nbr1, our previous work and this study show that its three ZZ domains are cargo-recognition modules (Liu et al., 2015). *S. cerevisiae* Atg19 does not have any ZZ domains. Instead, it uses three non-overlapping regions, the coiled-coil region (CC, residues 160-187), the Ape4-binding region (residues 204-247), and the Ams1 binding domain (ABD, residues 254-367), to bind Ape1, Ape4, and Ams1, respectively (Yuga et al., 2011; Watanabe et al., 2010; Yamasaki et al., 2016). The CC and the Ape4-binding region are not conserved in *S. pombe* Nbr1. The ABD in Atg19 has been proposed to be related to the functionally unknown FW domain (also called NBR1 domain) in human NBR1 and *S. pombe* Nbr1 (Kraft et al., 2010). Both the ABD in Atg19 and the FW domain in human NBR1 adopt an immunoglobulin-like beta-sandwich fold (Watanabe et al., 2010; Joint Center for Structural Genomics, 2014). The fact that *S. pombe* Nbr1 and *S. cerevisiae* Atg19 use completely different cargo-binding domains to recognize Ape4 and Ams1 suggests that cargo-receptor relationships can be maintained during evolution despite the loss of conservation in cargo-binding domains.

Unlike Atg19 in *S. cerevisiae*, syntenic orthologs of Atg19 in non-whole genome duplication (non-WGD) budding yeast species, as well as some post-WGD species, harbor ZZ domains (Figure S9A), suggesting that ZZ domains probably were lost in Atg19 proteins sometime after the WGD event occurring approximately 100 million years ago. In mammals, two autophagy receptors NBR1 and p62 both contain a ZZ domain. The function of the ZZ domain in NBR1 is unknown. The ZZ domain in p62 bind N-terminally arginylated proteins, which are considered not autophagy cargos, but rather activators of the autophagic activity of p62 (Zhang et al., 2018b; Ji et al., 2019).

This study, together with recently published structures of peptide-ligand-bound complexes of ZZ domains of p300, ZZZ3, p62, and HERC2 (Kwon et al., 2018; Zhang et al., 2018a; Mi et al., 2018; Zhang et al., 2018b; Liu et al., 2020), establish that a conserved mechanism of ZZ-mediated binding is the recognition of the N termini of ligand proteins by an acidic pocket on the ZZ domain (Figure S9B). This pocket contains three conserved residues, which are most often but not always aspartates (Figure S9C). This acidic pocket was likely present in the ancestor of Nbr1/Atg19-related proteins, as residues of this acidic pocket are conserved in Nbr1 homologs in the amoebozoan *Dictyostelium discoideum* and the basal metazoan *Nematostella vectensis* (sea anemone), as well as in ZZ-containing Atg19 orthologs (Figure S9C). ZZ2 and ZZ3 domains of Nbr1 have aspartates at all three positions of the acidic pocket (Figure S9C), consistent with our previous finding that the N-terminal residues of Ape2 is important for its vacuolar targeting (Liu et al., 2015). It is likely that recognizing the N-termini of cargos is a ubiquitous mechanism employed by ZZ-containing Nbr1/Atg19-family autophagy receptors.

At the atomic level, the conserved residues in the acidic pocket mainly interact with the N-terminal α-amino group and the backbone groups of the first peptide bond in the ligand (Figure S9B). As a consequence, ZZ domains sometime can tolerate variations of the N-terminal sequences of ligands, thereby allowing multi-specificity. Examples of multi-specificity include the ability of the ZZ domain of p62 to bind different types of N-degron peptides (Kwon et al., 2018), the binding of histone H3 and SUMO by the ZZ domain of HERC2 (Liu et al., 2020), and the binding of Ams1 and Ape4 by Nbr1-ZZ1. We note that, unlike Nbr1, in the cases of p62 and HERC2, the functional significance of the multi-specific binding ability of their ZZ domains is unclear. For Nbr1-ZZ1, N-terminal sequences of its ligands are not unconstrained, as evidenced by our mutational analysis results. The Ape4-ZZ1 complex structure shows that constraints on N-terminal sequences of ligands can be imposed by the requirement to maintain a proper conformation of the N-terminal tail through intramolecular interactions within the ligand protein.

Importantly, unlike other structures of ZZ-ligand complexes, in which the ligands are short peptides, our structures of ZZ1-cargo complexes contain full-length cargo proteins recognized by Nbr1-ZZ1. As a result, our structures reveal that ZZ domains can interact with non-N-terminal parts of partner proteins. This previously unknown binding mode can confer increased binding affinity, provide additional binding specificity, as well as allow more versatile ligand recognition. In the case of Nbr1-ZZ1, this binding mode acts cooperatively with the N-terminal binding mode to render possible the bispecific recognition of two distinct cargo proteins. It is likely that this binding mode is also used by other ZZ domains. Given that key residues for non-N-terminal binding mode of Nbr1-ZZ1, L66 and P97, are not conserved (Figure S9C), the versatility of the non-N-terminal binding mode is expected to be extensive and further studies will be needed to reveal the range of ligands recognized by ZZ domains.

## Methods

### Fission yeast strains and plasmids

Fission yeast strains used in this study are listed in Table S2 and plasmids used in this study are listed in Table S3. Genetic methods of strain construction and composition of media are as described (Forsburg and Rhind, 2006). The deletion strains used in this study were constructed by PCR amplifying the deletion cassettes in the Bioneer deletion strains and transforming PCR products into our laboratory strains or by standard PCR-based gene targeting. All point mutations were generated by PCR-based mutagenesis. Strains expressing proteins with different tags under native promoters were generated by PCR-based tagging. Plasmids overexpressing proteins with different tags under the control of *Pnmt1* or *P41nmt1* promoter were constructed using modified pDUAL vectors (Matsuyama et al., 2004). For the Pil1 co-tethering assay, truncated versions of Nbr1 were fused to the C terminus of Pil1-mCherry and placed under the control of *P41nmt1* promoter. The pDUAL plasmid expressing the Ams1-Nbr1 fusion protein was constructed by first using overlap-extension PCR to assemble together the following four fragments: the full-length Ams1, a sequence encoding a 13-residue linker (GFKKASSSDNKEQ), Nbr1(53-180), and maltose binding protein (MBP), and then cloning the PCR product upstream of two tandem human rhinovirus (HRV) 3C protease recognition sequence (LEVLFQGP) and GFP in a pDUAL vector. The plasmid expressing Ape4-Nbr1 fusion protein was constructed by first using overlap-extension PCR to assemble the following five fragments: the full-length Ape4, the same 13-residue linker described above, Nbr1(53-129), the same 13-residue linker described above but with altered codons, and MBP, and then cloning the PCR product upstream of two HRV 3C protease recognition sequences and GFP in a pDUAL vector. For the transformation of pDUAL plasmids into fission yeast, they were linearized with NotI digestion and integrated at the *leu1* locus, or linearized with MluI digestion and integrated at the *ars1* locus.

### Antibodies

The antibodies used for immunoblotting include: anti-Myc mouse monoclonal antibody (Abmart, Shanghai, China); peroxidase-anti-peroxidase for detecting TAP-tagged proteins (Sigma); anti-FLAG mouse monoclonal antibody (Sigma); anti-GST mouse monoclonal antibody (Abmart); anti-GFP mouse monoclonal antibody (Roche); anti-mCherry mouse monoclonal antibody (Abmart); rabbit polyclonal antibodies against Ape4, Ape2 and Lap2 (generated at NIBS antibody facility using recombinant Ape4, Ape2, and Lap2 as antigens).

### Affinity purification coupled with mass spectrometry (AP-MS) analysis

The bait proteins used were Nbr1 or Lap2 fused with a C-terminal YFP-FLAG-His_6_ tag and overexpressed from the *Pnmt1* promoter, or Ape2 fused with a C-terminal Venus tag and expressed from the endogenous promoter. The cell pellet was mixed with Lysis Buffer (50 mM HEPES-NaOH, pH 7.5, 150 mM NaCl, 1 mM EDTA, 1 mM dithiothreitol, 1 mM phenylmethylsulfonyl fluoride, 0.05% NP-40, 10% glycerol, 1×Roche protease inhibitor cocktail). Cell lysates were prepared by bead-beating. Bead beating lysis was performed using FastPrep-24 (MP Biomedicals). The lysates were incubated with GFP-Trap agarose beads (ChromoTek) for 3 h at 4°C. After incubation, the beads were washed 2 times using Lysis Buffer and another 2 times using Lysis Buffer without NP-40. Bead-bound proteins were eluted by incubation at 65°C with elution buffer (1% SDS, 100 mM Tris, pH 8.0).

Eluted proteins were precipitated with 20% TCA for at least 1 h. Protein precipitates were washed three times using ice-cold acetone and then dissolved in 8 M urea, 100 mM Tris, pH 8.5, reduced with 5 mM TCEP for 20 min, and alkylated with 10 mM iodoacetamide for 15 min in the dark. Then the samples were diluted by a factor of 4 and digested overnight at 37°C with trypsin (dissolved in 2 M urea, 1 mM CaCl_2_, 100 mM Tris, pH 8.5). Formic acid was added to the final concentration of 5% to stop digestion reaction. After digestion, the LC-MS/MS analysis was performed on an Easy-nLC II HPLC (Thermo Fisher Scientific) coupled to an LTQ Orbitrap XL mass spectrometer (Thermo Fisher Scientific). Peptides were loaded on a pre-column (100 μM ID, 4 cm long, packed with C12 10 μm 120 Å resin from YMC) and separated on an analytical column (75 μM ID, 10 cm long, packed with Luna C18 3 μm 100 Å resin from Phenomenex) using an acetonitrile gradient from 0–8% in 100 min at a flow rate of 200 nl/min. The top 8 most intense precursor ions from each full scan (resolution 60,000) were isolated for CID MS2 (normalized collision energy 35) with a dynamic exclusion time of 60 s. MS/MS fragment ions were detected by linear ion trap in a normal scan mode. Precursors with less than 2+ or unassigned charge states were excluded. The MS/MS spectra were searched with Prolucid against an *S. pombe* protein database. The search results were filtered with DTASelect.

### Immunoprecipitation

About 100 OD_600_ units of cells were collected and washed twice with water. The cell pellet was mixed with 100 μl of Lysis Buffer and 800 μl of 0.5-mm-diameter glass beads. Bead beating lysis was performed using FastPrep-24 (MP Biomedicals). Then another 300 μl of Lysis Buffer was added, and the cell lysate was cleared by centrifugation at 13000 rpm for 30 min. The supernatant was incubated with tag-recognizing antibody beads, including GFP-Trap agarose beads (ChromoTek) for mECitrine fusion proteins, RFP-Trap agarose beads (ChromoTek) for mCherry fusion proteins, and IgG Sepharose beads (GE Healthcare) for TAP-tagged proteins. After incubation, beads were washed 3 times with Lysis Buffer and eluted using SDS loading buffer.

### Fluorescence microscopy

Cells were grown to mid-log phase in EMM medium at 30°C. Microscopy was performed using the DeltaVision PersonalDV system (Applied Precision) equipped with a mCherry/CFP/YFP filter set (Chroma 89006 set) and a Photometrics Evolve 512 EMCCD camera. Images were obtained with a 100×1.4NA objective and analyzed with the SoftWoRx software.

### Immunoblotting assay examining vacuolar processing of mECitrine-tagged proteins

About 10 OD_600_ units of cells were harvested, mixed with 100 μl of Lysis Buffer, and lysed using the bead-beating lysis method. The lysate was mixed with 2×SDS loading buffer and boiled for 10 min. Samples were separated on a 12% SDS-PAGE gel and immunoblotted with anti-GFP antibody.

### Pil1 co-tethering assay

To examine a pair-wise protein-protein interaction, a bait protein was fused to Pil1-mCherry, and a prey protein was fused to mECitrine or CFP. Cells co-expressing both proteins were grown to log phase for fluorescence microscopy. To image the plasma-membrane associated filament-like structures formed by a Pil1-fused protein and its interactor, we acquired 8-10 optical Z-sections so that either the top or bottom plasma membrane is in focus in one of the Z-sections. Images were processed by deconvolution using the SoftWoRx software.

### Protein preparation for the structure determination

Fission yeast cells expressing the Ams1-Nbr1 fusion protein or the Ape4-Nbr1 fusion protein from the *Pnmt1* promoter were grown to mid-log phase in EMM medium at 30 °C. About 2000 OD_600_ units of cells were harvested and washed once with water. Cells were lysed by grinding in liquid nitrogen. The resulting powder was mixed with Lysis Buffer. After centrifugal clarification, the cell lysate was incubated with GFP-Trap Sepharose beads (ChromoTek) for 3 h at 4 °C. The beads were washed 4 times with wash buffer (50 mM HEPES-NaOH, pH 7.5, 150 mM NaCl, 1 mM EDTA, 1 mM dithiothreitol, 0.05% NP-40, 10% glycerol) and incubated with 100 μl of Lysis Buffer and 2 μg of 3C protease overnight at 4 °C. Protease-released proteins were concentrated and buffer-exchanged to storage buffer (50 mM Tris-HCl, pH 7.5, 150 mM NaCl, 5 mM MgCl_2_) by using an Amicon Ultra-0.5 centrifugal filter with 30 kDa molecular weight cutoff (Millipore).

### Ams1 and Ape4 competition assay

To examine whether Ams1 and Ape4 compete with each other for Nbr1 binding, we performed a GST pull down assay. GST-Nbr1(53-129) was purified from *E. coli* and FLAG_2_-His_6_ (FFH)-tagged Ams1 and Ape4 proteins were purified from *S. pombe*. Glutathione-Sepharose beads (GE Healthcare) preloaded with GST-Nbr1(53-129) were incubated with 50 μg Ams1-FFH, and then 25 μg, 100 μg, or 300 μg of Ape4-FFH was added to compete with Ams1-FFH for Nbr1 binding at 4°C for 3 h. Beads were washed 4 times with pull down buffer (50 mM NaH_2_PO_4_, pH 8.0, 300 mM NaCl, 10% glycerol) and eluted by SDS loading buffer. The eluates were separated by 12% SDS-PAGE gel and examined by immunoblotting using anti-FLAG antibody (for Ams1-FFH and Ape4-FFH), and anti-GST antibody (for GST-Nbr1(53-129)).

### Cryo-EM data collection

To prepare vitrified specimens, 3 μl of 0.5 mg/ml protein solutions was spread on a glow-charged holey carbon or NiTi film on Au grid and incubated for 10 sec in an FEI Vitrobot chamber set to 100% humidity and 4 °C. The grid was blotted for 3-5 sec and rapidly plunged into liquid ethane. Frozen grids were stored in liquid nitrogen.

The cryogenic samples were first examined on a Talos F200C 200kV electron microscope equipped with a Ceta camera (FEI). High-resolution data were collected on a 300kV Titan Krios electron microscope (FEI) equipped with a K2 Summit camera and a quantum energy filter (GIF, Gatan). The microscopy was operated in a zero-energy-loss mode with a slit width of 20 eV. Micrographs comprising of 36 frames each were collected using SerialEM (Mastronarde, 2005). The images were collected with a pixel size of 0.52 Å in the super-resolution counting mode and binned to a physical pixel size of 1.04 Å.

### Cryo-EM image processing

Movie stacks were subjected to motion correction with MotionCor2 (Zheng et al., 2017). Parameters of contrast transfer function (CTF) were estimated with Gctf (Zhang, 2016). Particles were automatically picked using gautomatch (http://www.mrclmb.cam.ac.uk/kzhang/Gautomatch/).

For the Ams1-Nbr1 complex, particles were extracted with a box size of 320 pixels and processed with a mask of 300 Å diameter. Particles were downsized 4-fold for 2D and 3D classification in RELION-3.0-beta (Zivanov et al., 2018). The free Ams1 structure (EMD-30021) low-pass filtered to 30 Å was used as the initial model for 3D classification and refinement. After one round of 2D classification and two rounds of 3D classification, 227,295 particles from high resolution classes were subjected for 3D reconstruction with D2 symmetry imposed in RELION-3.0-beta (Figure S4D). After the raw movie datasets were further processed in RELION for motion correction, particle polishing and CTF refinement, 3D refinement yielded a final density map of 2.4 Å resolution.

The Ape4-Nbr1 data were collected and processed similarly. Particles were extracted with a box size of 320 pixels and processed with a mask of 250 Å diameter. The crystal structure of human aspartyl aminopeptidase (PDB code 4DYO) was used as the initial model for 3D classification and refinement. After one round of 3D classification, 352,316 particles were subjected to 3D reconstruction with T symmetry imposed and the option of CTF refinement (Figure S6D). After post-processing in RELION, a density map of 2.26 Å resolution was obtained. The density map was also post-processed with LocScale (Jakobi et al., 2017), yielding a better resolved density for Nbr1-ZZ1 (Figure S7A).

### Model building and refinement

Structural models were built and adjusted in COOT (Emsley and Cowtan, 2004) and refined in real space using PHENIX with secondary structure, geometry, and noncrystallographic symmetry restrains (Adams et al., 2010). The statistics for data collection and structure refinement are summarized in Table S1.

To build the Ams1-Nbr1 complex structure, the model of Ams1 was derived from the cryo-EM structure of free Ams1 (PDB code 6LZ1) and the Nbr1-ZZ1 domain structure was de novo built. The model was refined against the sharpened cryo-EM map of Ams1-Nbr1 complex.

To build the Ape4-Nbr1 complex structure, a starting model of Ape4 was generated by Phyre2 on the template of hDNPEP structure (PDB code 3L6S). The structure of Nbr1-ZZ1 domain determined in complex of Ams1 was fitted as rigid body into the unsharpened and unfiltered cryo-EM map of the Ape4-Nbr1 complex. The final model of Ape4-Nbr1 complex was refined against the LocScale map.

The current model of Ams1-Nbr1 complex contains residues 2-1077 of Ams1, residues 1-3 of the linker and residues 59-108 of Nbr1. The current model of Ape4-Nbr1 complex contains residues 1-473 of Ape4, residues 1-2 of the linker and residues 59-108 of Nbr1.

Structural figures were prepared with PyMOL (Schrödinger, LLC) and Chimera (Pettersen et al., 2004). The inter-map FSC curve was calculated with relion_image_handle. To calculate FSC between model and map, the model was transformed into a density map in Chimera. The resultant model map was resampled to the grid of the map used for refinement. For cross-validation, the model was re-refined against a map reconstruction from the other half of the data. The resolution of density map was estimated using the criteria of gold-standard Fourier shell correlation (FSC) of 0.143 (Scheres and Chen, 2012). Local resolution was calculated with ResMap (Kucukelbir et al., 2014). Buried accessible surface area was calculated with areaimol using a probe of 1.4 Å radius (Vallone et al., 1998).

### 3D variability analysis

The distribution of Nbr1-ZZ1 in the Ape4-ZZ1 complex was assessed by 3D variability analysis (Punjani, 2020). 352,316 particles were exported to CryoSPARC 2.15 and subjected to non-uniform refinement (Punjani et al., 2017). These particles were expanded with T symmetry, resulting in 12-fold more particle images with each of the 12 subunits aligned to the same orientation. Subsequently, 3D variability analysis was performed with a mask covering one or two units of Ape4-ZZ1 complex, twenty clusters of volumes, and three eigenvectors. The results were visualized with the 3D Variability Display tool.

### Database

The cryo-EM density map and coordinates have been deposited to the Electron Microscopy Data Bank (EMDB) and Protein Data Bank (PDB) under accession numbers EMD-30650 and PDB 7DD9 for the Ams1-ZZ1 complex and EMD-30652 and PDB 7DDE for the Ape4-ZZ1 complex.

## Acknowledgements

We thank Boling Zhu, Xiaojun Huang, Gang Ji, Deyin Fan, Fei Sun and other staff at the Center for Biological Imaging, Institute of Biophysics, Chinese Academy of Sciences for their assistance with cryo-EM data collection. This work was supported by National Natural Science Foundation of China [32071199, 91940302], Strategic Priority Research Program of Chinese Academy of Sciences [XDB37010201], and National Key R&D Program of China [2017YFA0504600] to K.Y. and grants from the Chinese Ministry of Science and Technology and the Beijing municipal government to M.-Q.D. and L.-L.D.

## Author contributions

Conceptualization: Y.-Y.W., J.Z., K.Y., and L.-L.D.; Methodology and investigation: Y.-Y.W., J.Z., X.-M.L., M.-Q.D., K.Y., and L.-L.D.; Writing – original draft: Y.-Y.W. and J.Z.; Writing – review and editing: K.Y. and L.-L.D.; Funding acquisition: K.Y., M.-Q.D., and L.-L.D.

---

**Supplementary figure 1.**
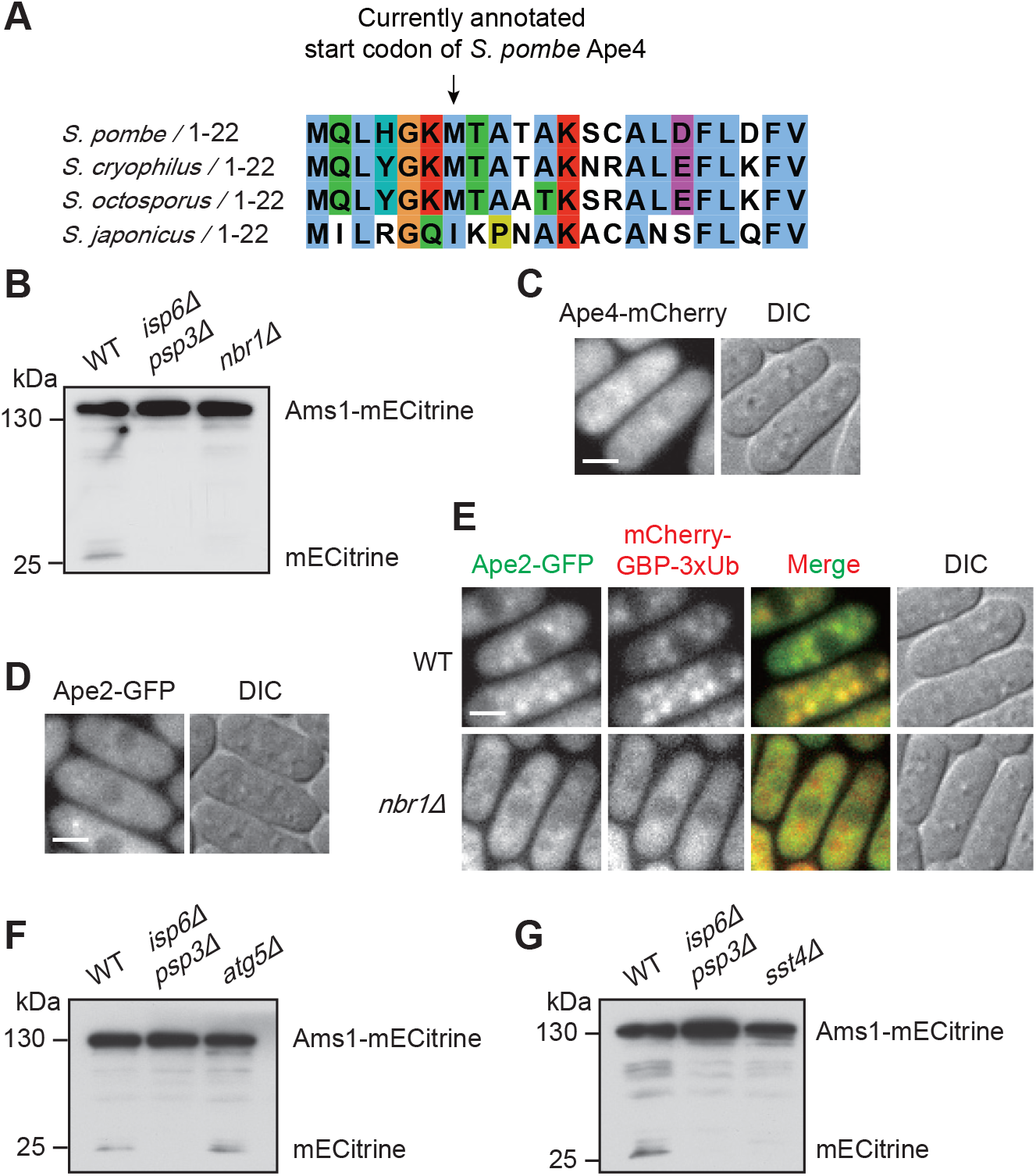
Vacuolar targeting of Ams1 and Ape4 requires Nbr1 and ESCRTs, but not Atg proteins. (A) Sequence alignment of the N-terminal region of Ape4 proteins from four fission yeast species. (B) Ams1-mECitrine was processed to free mECitrine in a manner that depends on vacuolar proteases Isp6 and Psp3 and also depends on Nbr1. Vacuolar processing of Ams1-mECitrine was examined by immunoblotting (IB). (C) Localization of Ape4. Ape4 was endogenously tagged at its C-terminus with mCherry. Bar, 3 μm. (D) Ape2 endogenously tagged at its C-terminus with GFP did not exhibit obvious vacuole localization. Bar, 3 μm. (E) Vacuolar targeting of Ape2-GFP was enhanced by tethering 3xUb to it. mCherry-GBP-3xUb was expressed under the *P41nmt1* promoter. Bar, 3 μm. (F) Vacuolar processing of Ams1-mECitrine was not affected by *atg5Δ*. (G) Vacuolar processing of Ams1-mECitrine was abolished by *sst4Δ*.

**Supplementary figure 2.**
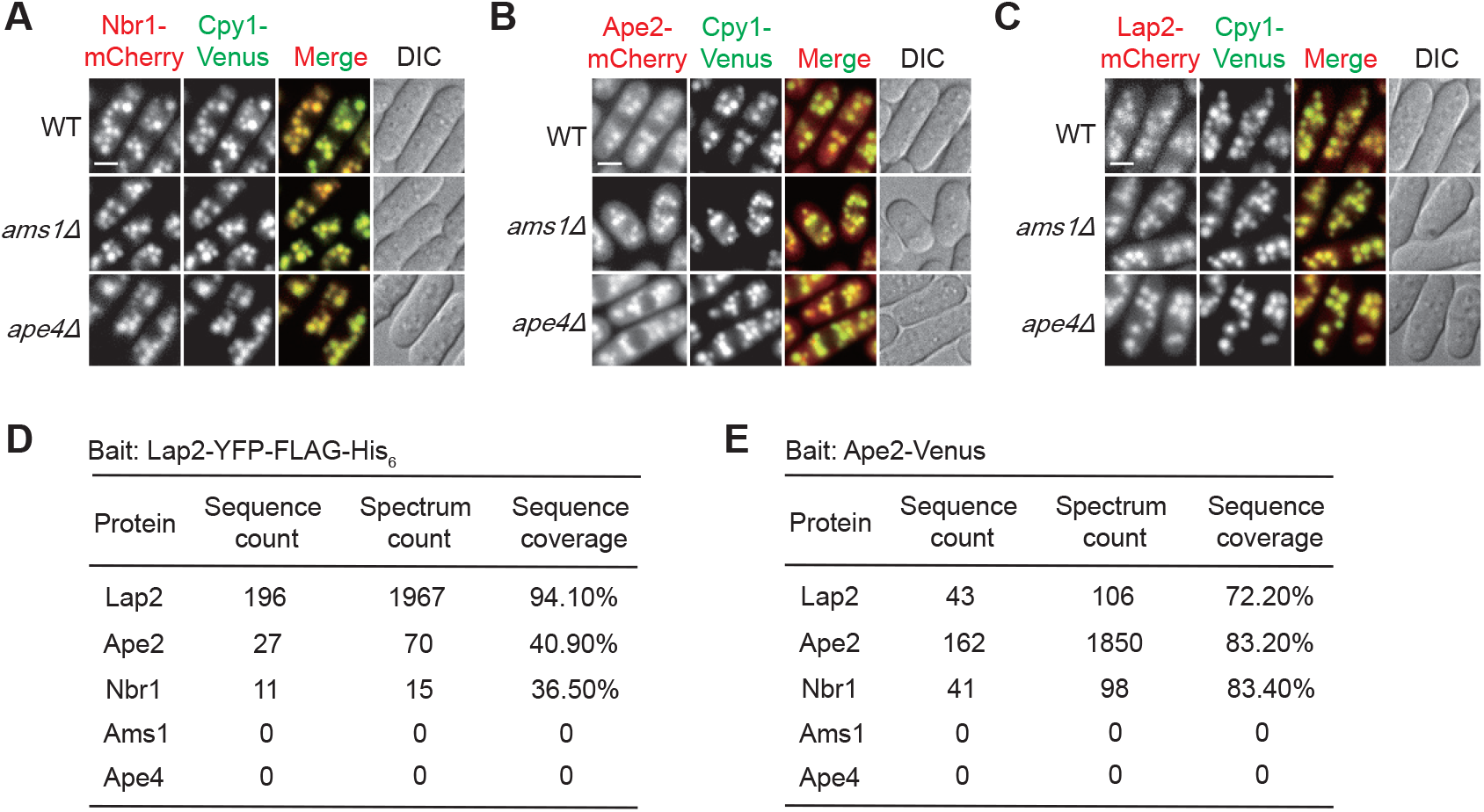
Ams1 and Ape4 are not required for vacuolar targeting of Nbr1, Ape2, and Lap2 and do not interact with Lap2 or Ape2. (A) Localization of Nbr1 in WT, *ams1Δ*, and *ape4Δ* cells. Bar, 3 μm. (B) Localization of Ape2 in WT, *ams1Δ*, and *ape4Δ* cells. Bar, 3 μm. (C) Localization of Lap2 in WT, *ams1Δ*, and *ape4Δ* cells. Bar, 3 μm. (D) In an AP-MS analysis using Lap2 as bait, Nbr1 and Ape2, but not Ams1 and Ape4 co-purified with Lap2. (E) In an AP-MS analysis using Ape2 as bait, Nbr1 and Lap2, but not Ams1 and Ape4. co-purified with Ape2.

**Supplementary figure 3.**
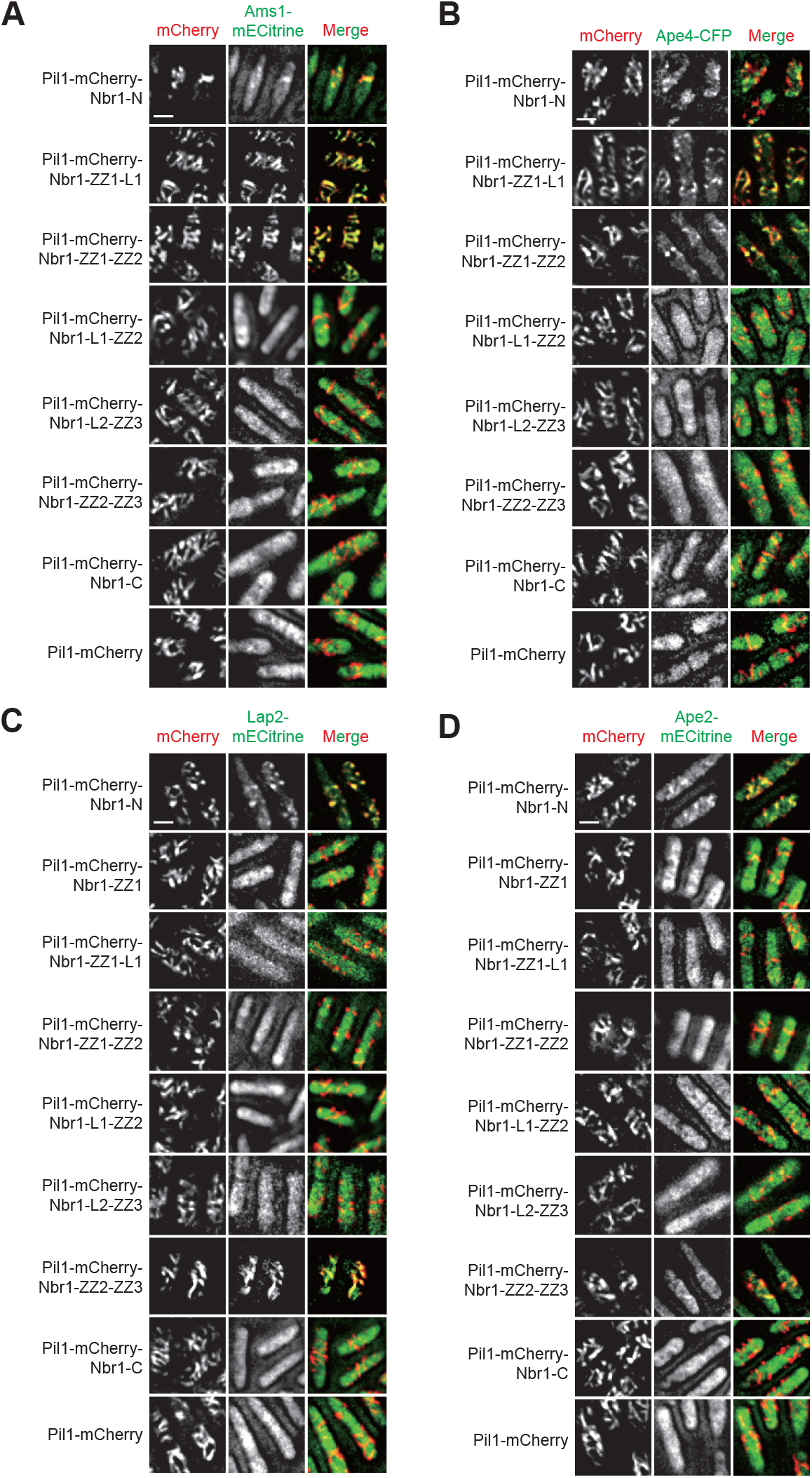
Truncation analysis of Nbr1. (A) The Pil1 co-tethering assay was used to determine whether truncated fragments of Nbr1 can interact with Ams1. Truncated Nbr1 fragments were fused to the C-terminus of Pil1-mCherry. The strain expressing Pil1-mCherry served as a negative control. Bar, 3 μm. (B) The Pil1 co-tethering assay was used to determine whether truncated fragments of Nbr1 can interact with Ape4. Bar, 3 μm. (C) The Pil1 co-tethering assay was used to determine whether truncated fragments of Nbr1 can interact with Lap2. Bar, 3 μm. (D) The Pil1 co-tethering assay was used to determine whether truncated fragments of Nbr1 can interact with Ape2. Bar, 3 μm.

**Supplementary figure 4.**
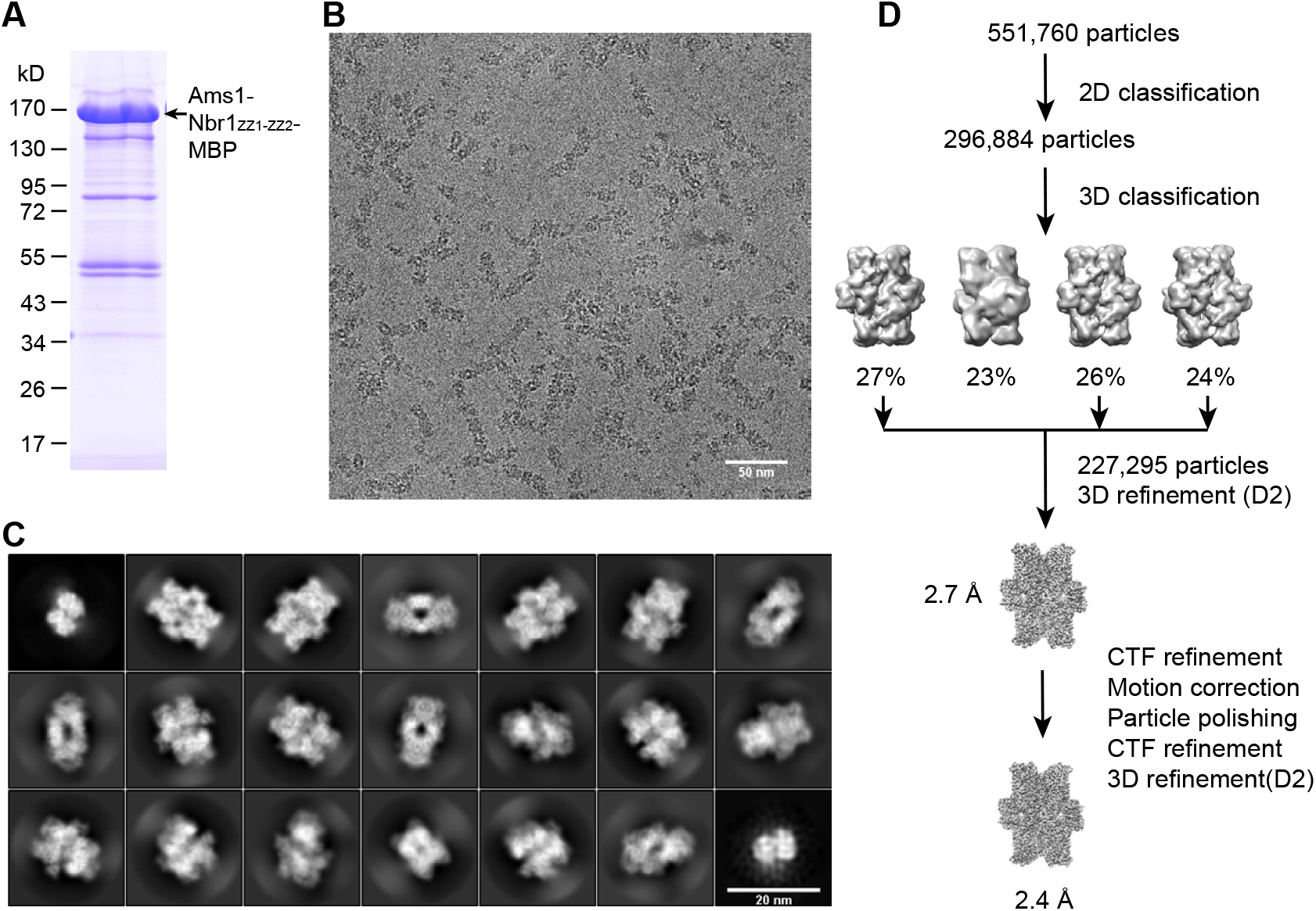
Cryo-EM analysis of the Ams1-ZZ1 complex. (A) An SDS-PAGE analysis of the purified Ams1-Nbr1_ZZ1-ZZ2_-MBP fusion protein used for cryo-EM analysis. (B) A typical cryo-EM micrograph. Bar, 50 nm. (C) Typical 2D class averages from reference-free alignment. Bar, 20 nm. (D) Flowchart of 2D classification, 3D classification, and refinement.

**Supplementary figure 5.**
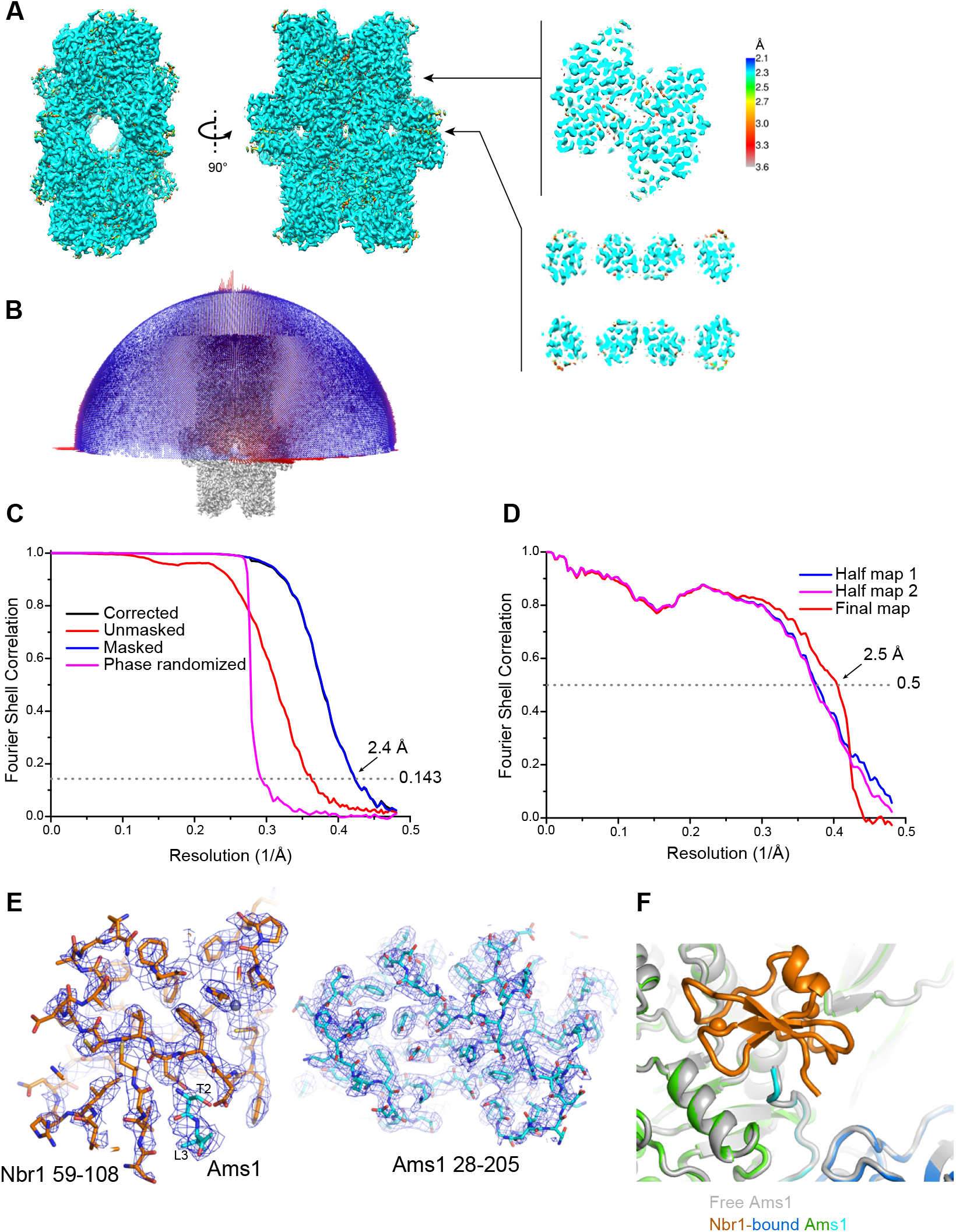
Quality of the cryo-EM map of the Ams1-ZZ1 complex and superposition of structures of Ams1. (A) Cryo-EM density map of the Ams1-ZZ1 complex colored by local resolution. Two orthogonal views of the surface are displayed. Two cut-through views are shown on the right. (B) Angular distribution of particles. Each cylinder represents one orientation with height proportional to number of particles. (C) FSC curves for the corrected (black), unmasked (red), masked (blue), and phase-randomized (magenta) maps. The overall resolution is 2.4 Å according to the FSC=0.143 criterion. (D) Cross-validation FSC curves of the atomic model versus map. Blue: FSC between the re-refined model and half-map 1 against which the model was refined. Magenta: FSC between the re-refined model and half-map 2 against which the model was not refined. Red: FSC between the refined structure and the final post-processed map. (E) Representative cryo-EM densities with superimposed structural models. (F) Structural superposition of free and ZZ1-bound Ams1. The structure of free Ams1 (PDB code 6LZ1) is colored in grey.

**Supplementary figure 6.**
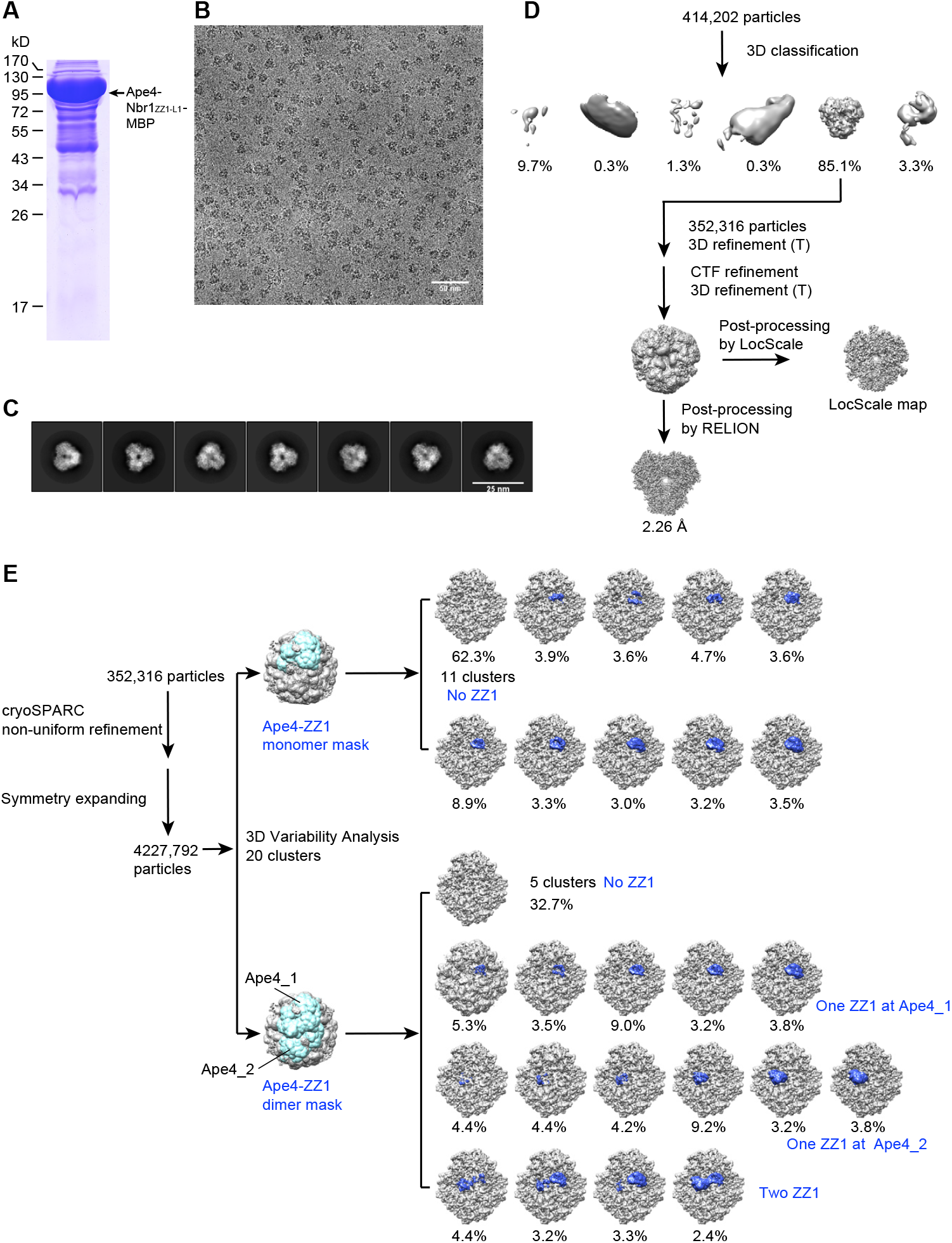
Cryo-EM analysis of the Ape4-ZZ1 complex. (A) An SDS-PAGE analysis of purified Ape4-Nbr1_ZZ1-ZZ2_-MBP protein used for cryo-EM analysis. (B) A typical cryo-EM micrograph. Bar, 50 nm. (C) Typical 2D class averages from reference-free alignment. Bar, 25 nm. (D) Flowchart of 3D classification and refinement. (E) 3D variability analysis of Nbr1-ZZ1 in the Ape4-ZZ1 complex. 20 clusters of volumes are classified according to a mask covering one or two Ape4-ZZ1 complexes and sorted by Nbr1-ZZ1 occupancy. The density of Nbr1-ZZ1 is colored dark blue. The clusters without ZZ1 density were combined. Fractions of particles belonging to each cluster are indicated.

**Supplementary figure 7.**
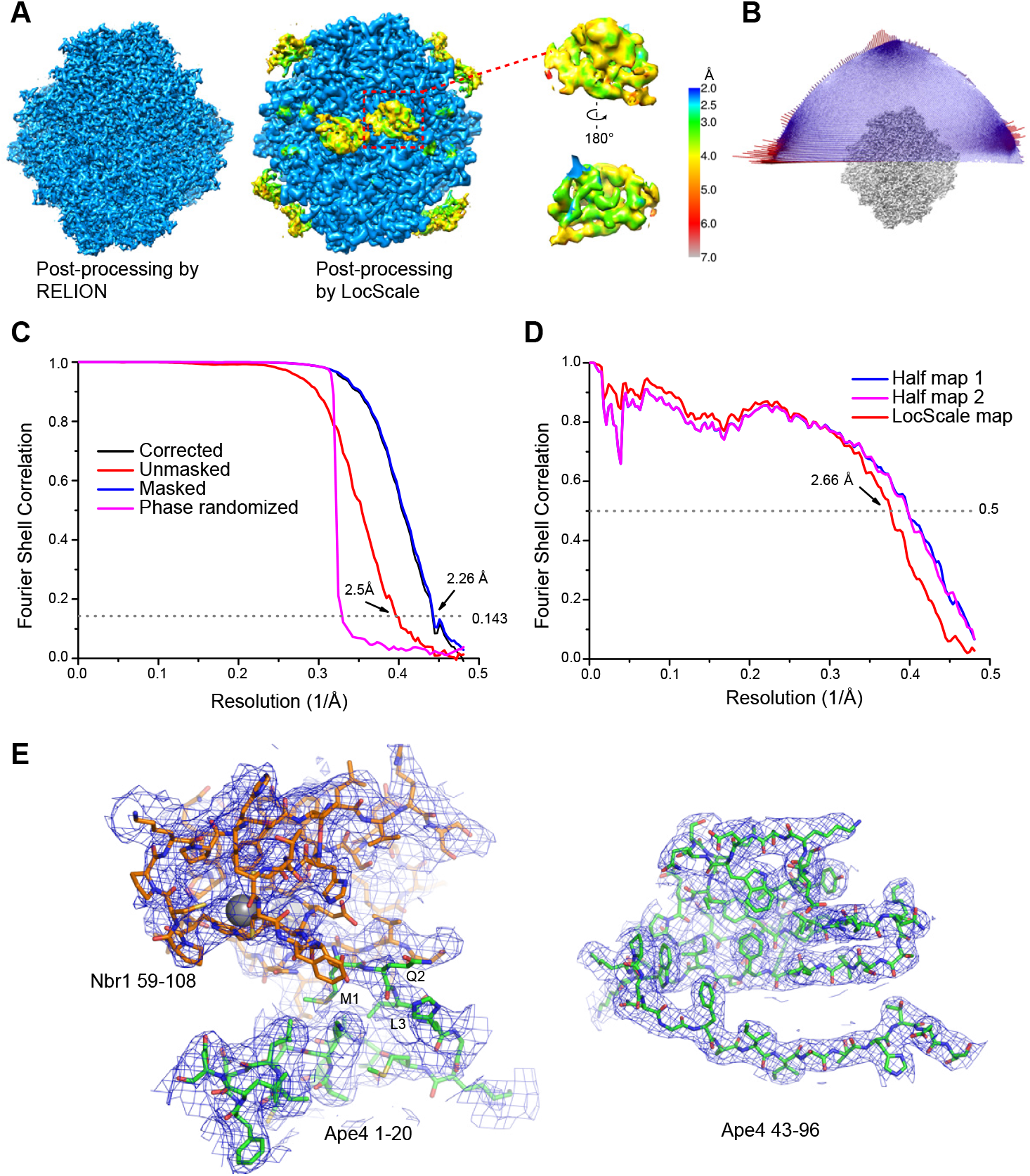
Quality of the cryo-EM map of the Ape4-ZZ1 complex. (A) Cryo-EM density maps of the Ape4-ZZ1 complex colored by local resolution. The map sharpened by RELION is shown on the left where Nbr1 is invisible. The LocScale map is shown on the right with visible Nbr1. The density of Nbr1-ZZ1 is also extracted and shown in two opposite orientations. (B) Angular distribution of particles. Each cylinder represents one orientation with height proportional to number of particles. (C) FSC curves for the corrected (black), unmasked (red), masked (blue), and phase-randomized (magenta) maps. The overall resolution is 2.26 Å according to the FSC=0.143 criterion. (D) Cross-validation FSC curves of the atomic model versus map. Blue: FSC between the re-refined model and half-map 1 against which the model was refined. Magenta: FSC between the re-refined model and half-map 2 against which the model was not refined. Red: FSC between the refined structure and LocScale map. (E) Representative cryo-EM densities from the LocScale map with superimposed structural models.

**Supplementary figure 8.**
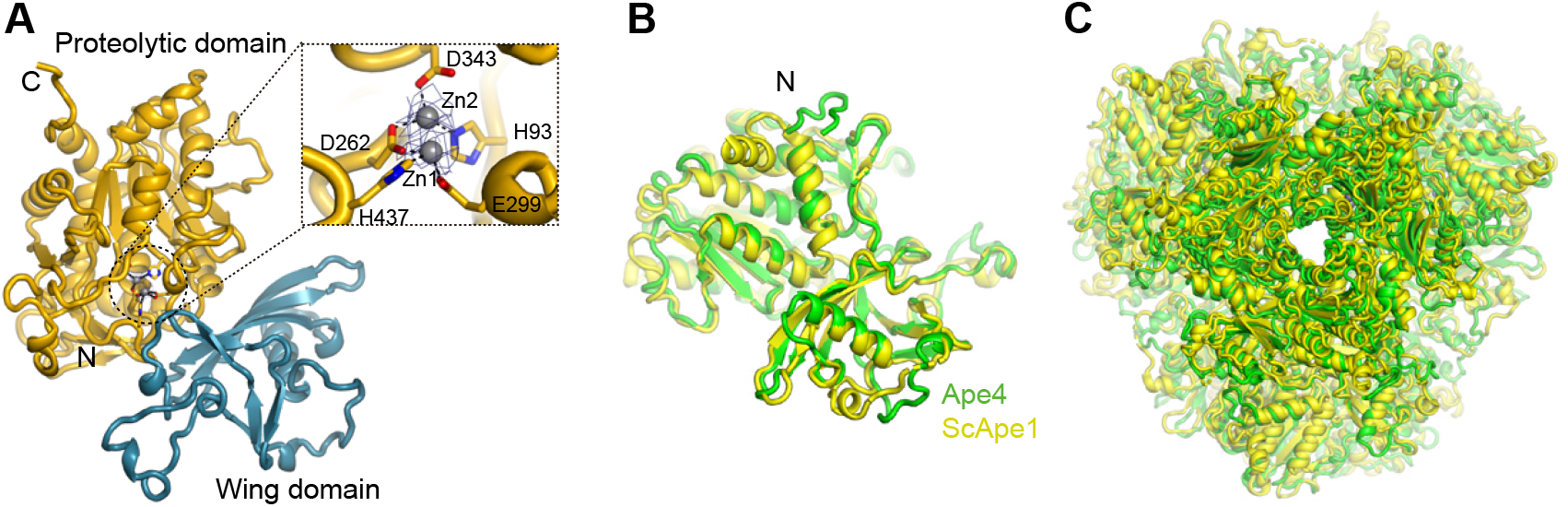
Structure of the Ape4-ZZ1 complex. (A) Structure of an Ape4 monomer. The wing domain and the proteolytic domain are colored in blue and orange, respectively. The insert shows the binding of two zinc ions (spheres) in the active site, with coordinating bonds shown as dashed lines. Densities around the zinc ions are displayed as mesh. (B) Superposition of the monomer structures of Ape4 (green) and the precursor form of *Saccharomyces cerevisiae* Ape1 (ScApe1) (yellow) (PDB code 5JH9). The N-termini are labeled. The RMSD is 0.82 Å over 352 Cα atom pairs. (C) Superposition of the dodecamer structures of Ape4 and ScApe1.

**Supplementary figure 9.**
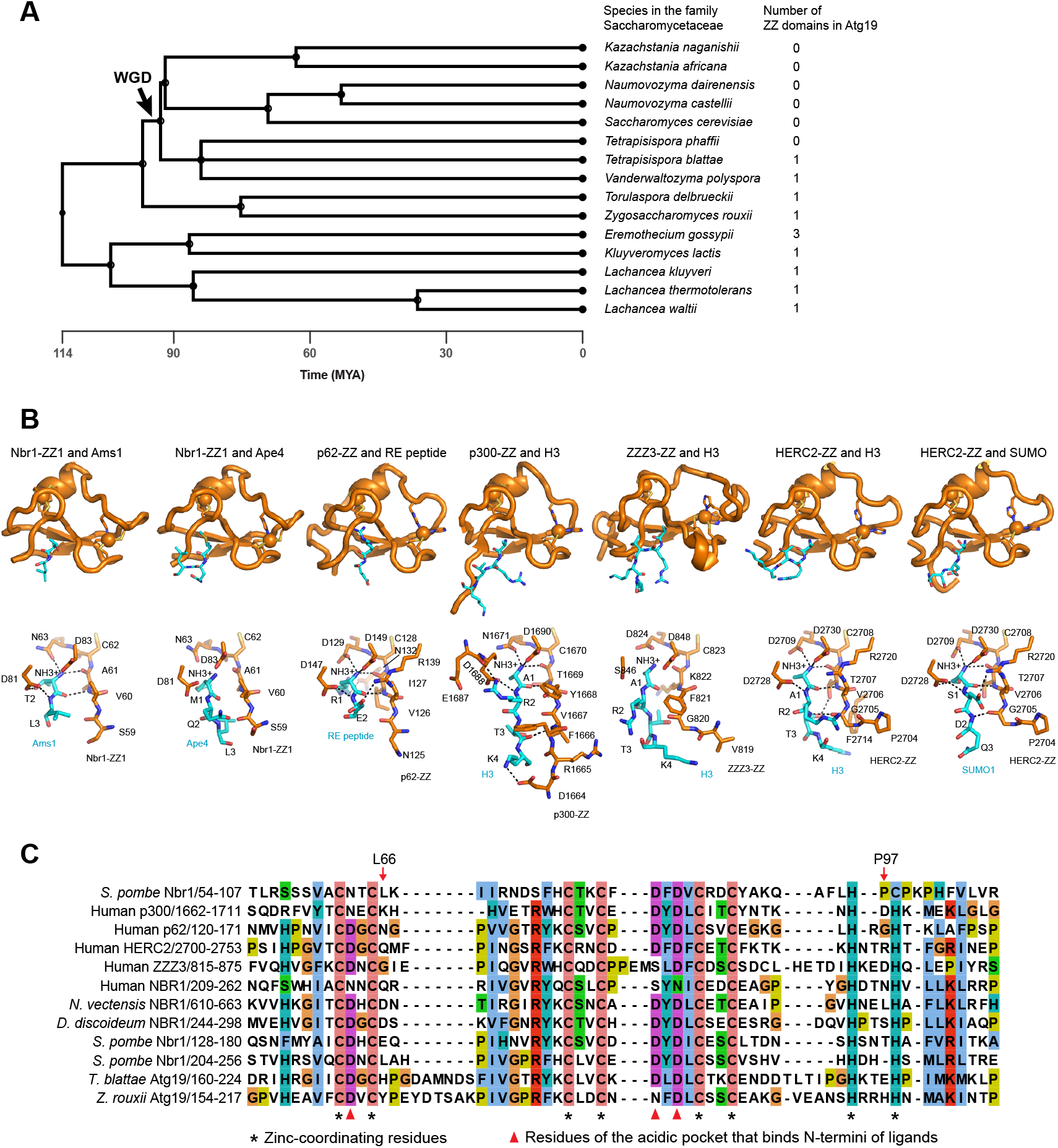
Dating the loss of ZZ domains in Atg19 proteins, structural comparison of ZZ-ligand complexes, and sequence alignment of ZZ domains. (A) A timetree of 15 Saccharomycetaceae species and the number of ZZ domains in Atg19 orthologs in these species. The timetree was generated using the TimeTree resource (http://www.timetree.org)(Kumar et al., 2017). Sequences of syntenic orthologs of *S. cerevisiae* Atg19 were retrieved using the Yeast Gene Order Browser (YGOB; http://ygob.ucd.ie)(Byrne and Wolfe, 2005). (B) Structural comparison highlighting the N-terminal binding mode of ZZ domains. The aligned ZZ domain structures are from Nbr1 in complex with Ams1 or Ape4, p62 bound to an N-degron dipeptide Arg-Glu (PDB code 6MIU), p300 bound to the N-terminal peptide of histone H3 (PDB code 6DS6), ZZZ3 bound to the N-terminal peptide of histone H3 (PDB code 6E83), and HERC2 bound to the N-terminal peptide of histone H3 (PDB code 6WW4) or the N-terminal peptide of SUMO (PDB code 6WW3). The overall structure (top) and detailed interactions with the bound peptides (bottom) are shown for each ZZ-peptide complex. (C) Sequence alignment of ZZ domains. The top five sequences correspond to the ZZ domains whose structures are compared in (B). The residues of Nbr1-ZZ1 involved in binding non-N-terminal parts of cargos, L66 and P97, are highlighted by red arrows. The accession numbers of the sequences are: Q9P792 (*S. pombe* Nbr1), Q09472 (p300), Q13501 (p62), O95714 (HERC2), Q8IYH5 (ZZZ3), NP_114068 (human NBR1), XP_001634545.1 (*Nematostella vectensis* NBR1), XP_646538.1 (*Dictyostelium discoideum* NBR1), XP_004181174.1 (*Tetrapisispora blattae* Atg19), and XP_002495208.1 (*Zygosaccharomyces rouxii* Atg19).

**Table S1.**
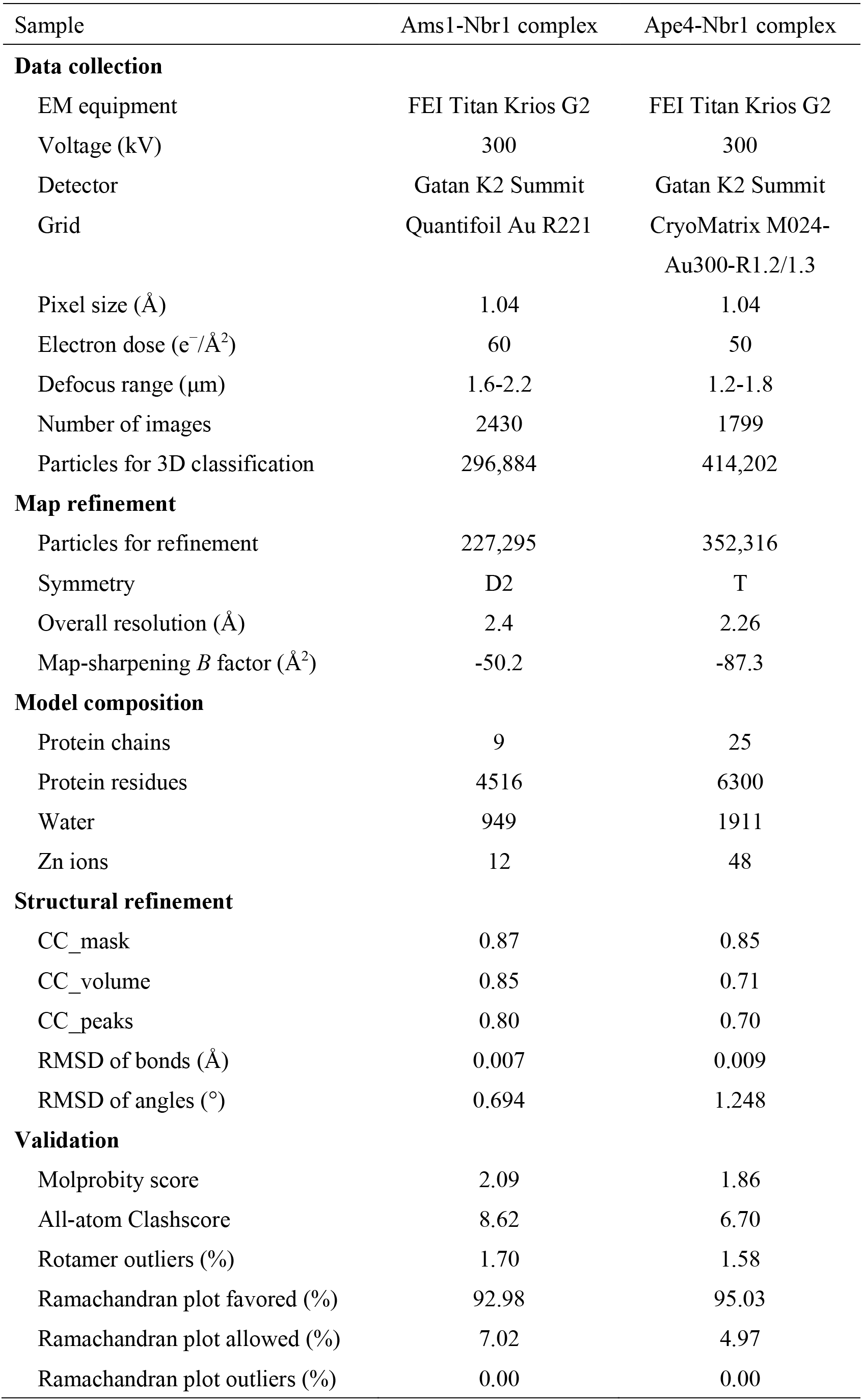
Statistics of data collection, structural refinement and model validation

