## Supplementary material for "Molecular and structural mechanisms of ZZ domain-mediated cargo recognition by autophagy receptor Nbr1": Tables S2 and S3

Table S2. Fission yeast strains used in this study.

| Strain | Mating type | Genotype |
|---|---|---|
| LD328 | h+ | *leu1-32 his3-D1* |
| DY6142 | h- | *leu1-32 his3-D1* |
| LD259 | h+ | *leu1-32 his3-D1 ura4-D18* |
| LD260 | h- | *leu1-32 his3-D1 ura4-D18* |
| DY26121 | h+ | *his3-D1 leu1-32::Pnmt1-nbr1-YFH(leu1+)* |
| xm765 | | *his3-D1 leu1-32::Pnbr1-nbr1-TAP(leu+) ams1-13Myc::KanMx nbr1Δ::HphMX* |
| xm708 | h- | *his3-D1 leu1-32::Pnbr1-nbr1-TAP(leu+) nbr1Δ::HphMX* |
| xm659 | h+ | *his3-D1 leu1-32 ams1-13Myc::KanMx* |
| wyy1236 | | *his3-D1 leu1-32::Pnbr1-nbr1-TAP(leu+) ape4-13Myc::KanMx* |
| xm667 | h+ | *his3-D1 leu1-32 ape4-13Myc::KanMx* |
| xm978 | h+ | *leu1-32 his3-D1 ams1-mCherry::KanMx cpy1-Venus::NatMx* |
| xm2344 | | *leu1-32 his3-D1 ams1-mCherry::KanMx cpy1-Venus::NatMx nbr1Δ::HphMX* |
| wyy2690 | | *leu1-32 his3-D1 ams1-mCherry::KanMx cpy1-Venus::NatMx sst4Δ::KanMX* |
| wyy584 | | *leu1-32 his3-D1 ams1-mCherry::KanMx cpy1-Venus::NatMx atg1Δ::KanMX* |
| wyy478 | h+ | *his3-D1 ura4-D18 leu1::P41nmt1-ams1-mECitrine(leu+)* |
| wyy518 | | *his3-D1 ura4-D18 leu1::P41nmt1-ams1-mECitrine(leu+) isp6Δ::HphMX psp3Δ::KanMX* |
| wyy512 | h+ | *his3-D1 ura4-D18 leu1::P41nmt1-ams1-mECitrine(leu+) nbr1Δ::KanMX* |
| wyy515 | | *his3-D1 ura4-D18 leu1::P41nmt1-ams1-mECitrine(leu+) sst4Δ::KanMX* |
| wyy503 | | *his3-D1 ura4-D18 leu1::P41nmt1-ams1-mECitrine(leu+) atg5Δ::NatMX* |
| xm618 | h- | *his3-D1 leu1-32 ape4-mCherry::KanMX* |
| xm460 | h+ | *his3-D1 leu1-32 ape2-GFP::KanMX* |
| wyy1179 | h+ | *his3-D1 leu1::P41nmt1-mCherry-GBP-3xUb(leu+) ape2-GFP::KanMX* |
| wyy1260 | | *his3-D1 leu1::P41nmt1-mCherry-GBP-3xUb(leu+) ape2-GFP::KanMX nbr1Δ::KanMX* |
| wyy2638 | | *his3-D1 leu1::P41nmt1-GBP-3xUb(leu+) ape4-GFP::KanMX cpy1-Venus::NatMX* |
| wyy2642 | | *his3-D1 leu1::P41nmt1-GBP-3xUb(leu+) ape4-GFP::KanMX cpy1-Venus::NatMX nbr1Δ::KanMX* |
| wyy2644 | | *his3-D1 leu1::P41nmt1-GBP-3xUb(leu+) ape4-GFP::KanMX cpy1-Venus::NatMX sst4Δ::KanMX* |
| wyy2640 | | *his3-D1 leu1::P41nmt1-GBP-3xUb(leu+) ape4-GFP::KanMX cpy1-Venus::NatMX atg1Δ::KanMX* |
| wyy572 | | *leu1-32 his3-D1 ams1-mCherry::KanMx cpy1-Venus::NatMx lap2Δ::HphMX* |

| | | |
|---|---|---|
| wyy573 | | *leu1-32 his3-D1 ams1-mCherry::KanMx cpy1-Venus::NatMx ape2Δ::HphMX* |
| wyy2691 | | *leu1-32 his3-D1 ams1-mCherry::KanMx cpy1-Venus::NatMx ape4Δ::KanMx* |
| wyy2729 | | *his3-D1 leu1::P41nmt1-GBP-3xUb(leu+) ape4-GFP::KanMX cpy1-Venus::NatMX lap2Δ::NatMX* |
| wyy2643 | | *his3-D1 leu1::P41nmt1-GBP-3xUb(leu+) ape4-GFP::KanMX cpy1-Venus::NatMX ape2Δ::KanMX* |
| wyy2726 | | *his3-D1 leu1::P41nmt1-GBP-3xUb(leu+) ape4-GFP::KanMX cpy1-Venus::NatMX ams1Δ::KanMX* |
| xm478 | h+ | *his3-D1 leu1-32 lap2-TAP::KanMx* |
| wyy525 | | *his3-D1 leu1-32 ams1-13Myc::HphMx lap2-TAP::KanMx* |
| xm661 | h- | *his3-D1 leu1-32 ams1-13Myc::HphMx* |
| xm738 | h+ | *his3-D1 leu1::Pnbr1-nbr1-TAP(leu+) ams1-mCherry::KanMx nbr1Δ::HphMX* |
| xm636 | h+ | *his3-D1 leu1::Pnbr1-nbr1-TAP(leu+) nbr1Δ::HphMX* |
| xm626 | h- | *his3-D1 leu1-32 ams1-mCherry::KanMx* |
| wyy1242 | | *his3-D1 leu1-32 ape4-TAP::HphMx ams1-mCherry::KanMx* |
| xm655 | h+ | *his3-D1 leu1-32 ape4-TAP::HphMx* |
| xm985 | h+ | *leu1-32 his3-D1 nbr1-mCherry::natMX cpy1-Venus::hphMX* |
| wyy1005 | | *leu1-32 his3-D1 nbr1-mCherry::natMX cpy1-Venus::hphMX ams1Δ::HphMX* |
| wyy1596 | | *leu1-32 his3-D1 nbr1-mCherry::natMX cpy1-Venus::hphMX ape4Δ::KanMX* |
| xm982 | h+ | *leu1-32 his3-D1 ape2-mCherry::kanMX cpy1-Venus::natMX* |
| wyy678 | | *leu1-32 his3-D1 ape2-mCherry::kanMX cpy1-Venus::natMX ams1Δ::HphMX* |
| wyy1651 | | *leu1-32 his3-D1 ape2-mCherry::kanMX cpy1-Venus::natMX ape4Δ::KanMX* |
| xm989 | h- | *leu1-32 his3-D1 lap2-mCherry::kanMX cpy1-Venus::natMX* |
| wyy1010 | | *leu1-32 his3-D1 lap2-mCherry::kanMX cpy1-Venus::natMX ams1Δ::HphMX* |
| wyy1602 | | *leu1-32 his3-D1 lap2-mCherry::kanMX cpy1-Venus::natMX ape4Δ::HphMX* |
| xm114 | h+ | *his3-D1 leu1::Pnmt1-lap2-YFH(leu+)* |
| xm464 | h+ | *leu1-32 his3-D1 ape2-Venus::kanMX* |
| wyy565 | h+ | *his3-D1 ars1::p41nmt1-pil1-mCherry-nbr1-N(ura+) leu1::P41nmt1-ams1-mECitrine(leu+)* |
| wyy2554 | h+ | *his3-D1 ars1::p41nmt1-pil1-mCherry-nbr1-ZZ1(ura+) leu1::P41nmt1-ams1-mECitrine(leu+)* |
| wyy797 | h+ | *his3-D1 ars1::p41nmt1-pil1-mCherry-nbr1-ZZ1-ZZ2(ura+) leu1::P41nmt1-ams1-mECitrine(leu+)* |
| wyy872 | h+ | *his3-D1 ars1::p41nmt1-pil1-mCherry-nbr1-ZZ2-ZZ3(ura+) leu1::P41nmt1-ams1-mECitrine(leu+)* |
| wyy1037 | h+ | *his3-D1 ars1::p41nmt1-pil1-mCherry-nbr1-ZZ1-L1(ura+) leu1::P41nmt1-ams1-mECitrine(leu+)* |
| wyy1080 | h+ | *his3-D1 ars1::p41nmt1-pil1-mCherry-nbr1-L1-ZZ2(ura+) leu1::P41nmt1-ams1-mECitrine(leu+)* |

| | | |
|---|---|---|
| wyy1041 | h+ | *his3-D1 ars1::p41nmt1-pil1-mCherry-nbr1-L2-ZZ3(ura+) leu1::P41nmt1-ams1-mECitrine(leu+)* |
| wyy2739 | h+ | *his3-D1 ars1::P41nmt1-ams1-mECitrine(ura+) leu1::p41nmt1-pil1-mCherry-nbr1-C(leu+)* |
| wyy817 | h+ | *his3-D1 ars1::p41nmt1-pil1-mCherry(ura+) leu1::P41nmt1-ams1-mECitrine(leu+)* |
| wyy1138 | | *his3-D1 ura? ape4-CFP::KanMx leu1::p41nmt1-pil1-mCherry-nbr1-N(leu+)* |
| wyy2551 | h- | *his3-D1 ape4-CFP::KanMx leu1::p41nmt1-pil1-mCherry-nbr1-ZZ1(leu+)* |
| wyy1200 | h- | *his3-D1 ape4-CFP::KanMx leu1::p41nmt1-pil1-mCherry-nbr1-ZZ1-ZZ2(leu+)* |
| wyy1214 | h- | *his3-D1 ape4-CFP::KanMx leu1::p41nmt1-pil1-mCherry-nbr1-ZZ2-ZZ3(leu+)* |
| wyy1197 | h- | *his3-D1 ape4-CFP::KanMx leu1::p41nmt1-pil1-mCherry-nbr1-ZZ1-L1(leu+)* |
| wyy1223 | h- | *his3-D1 ape4-CFP::KanMx leu1::p41nmt1-pil1-mCherry-nbr1-L1-ZZ2(leu+)* |
| wyy1194 | h- | *his3-D1 ape4-CFP::KanMx leu1::p41nmt1-pil1-mCherry-nbr1-L2-ZZ3(leu+)* |
| wyy1144 | | *his3-D1 ura? ape4-CFP::KanMx leu1::p41nmt1-pil1-mCherry-nbr1-C(leu+)* |
| wyy2536 | h- | *his3-D1 ape4-CFP::KanMx leu1::p41nmt1-pil1-mCherry(leu+)* |
| wyy777 | h+ | *his3-D1 ars1::p41nmt1-pil1-mCherry-nbr1-N(ura+) leu1::P41nmt1-lap2-mECitrine(leu+)* |
| wyy2558 | h+ | *his3-D1 ars1::p41nmt1-pil1-mCherry-nbr1-ZZ1(ura+) leu1::P41nmt1-lap2-mECitrine(leu+)* |
| wyy1020 | h+ | *his3-D1 ars1::p41nmt1-pil1-mCherry-nbr1-ZZ1-ZZ2(ura+) leu1::P41nmt1-lap2-mECitrine(leu+)* |
| wyy867 | h+ | *his3-D1 ars1::p41nmt1-pil1-mCherry-nbr1-ZZ2-ZZ3(ura+) leu1::P41nmt1-lap2-mECitrine(leu+)* |
| wyy1045 | h+ | *his3-D1 ars1::p41nmt1-pil1-mCherry-nbr1-ZZ1-L1(ura+) leu1::P41nmt1-lap2-mECitrine(leu+)* |
| wyy1085 | h+ | *his3-D1 ars1::p41nmt1-pil1-mCherry-nbr1-L1-ZZ2(ura+) leu1::P41nmt1-lap2-mECitrine(leu+)* |
| wyy1049 | h+ | *his3-D1 ars1::p41nmt1-pil1-mCherry-nbr1-L2-ZZ3(ura+) leu1::P41nmt1-lap2-mECitrine(leu+)* |
| wyy2695 | h+ | *his3-D1 ars1::P41nmt1-lap2-mECitrine(ura+) leu1::p41nmt1-pil1-mCherry-nbr1-C(leu+)* |
| wyy811 | h+ | *his3-D1 ars1::p41nmt1-pil1-mCherry(ura+) leu1::P41nmt1-lap2-mECitrine(leu+)* |
| wyy780 | h+ | *his3-D1 ars1::p41nmt1-pil1-mCherry-nbr1-N(ura+) leu1::P41nmt1-ape2-mECitrine(leu+)* |
| wyy818 | h+ | *his3-D1 ars1::p41nmt1-pil1-mCherry-nbr1-ZZ1(ura+) leu1::P41nmt1-ape2-mECitrine(leu+)* |
| wyy823 | h+ | *his3-D1 ars1::p41nmt1-pil1-mCherry-nbr1-ZZ1-ZZ2(ura+) leu1::P41nmt1-ape2-mECitrine(leu+)* |
| wyy868 | h+ | *his3-D1 ars1::p41nmt1-pil1-mCherry-nbr1-ZZ2-ZZ3(ura+) leu1::P41nmt1-ape2-mECitrine(leu+)* |
| wyy1051 | h+ | *his3-D1 ars1::p41nmt1-pil1-mCherry-nbr1-ZZ1-L1(ura+) leu1::P41nmt1-ape2-mECitrine(leu+)* |
| wyy1053 | h+ | *his3-D1 ars1::p41nmt1-pil1-mCherry-nbr1-L1-ZZ2(ura+) leu1::P41nmt1-ape2-mECitrine(leu+)* |
| wyy1058 | h+ | *his3-D1 ars1::p41nmt1-pil1-mCherry-nbr1-L2-ZZ3(ura+) leu1::P41nmt1-ape2-mECitrine(leu+)* |

| | | |
|---|---|---|
| wyy2697 | h+ | *his3-D1 ars1::P41nmt1-ape2-mECitrine(ura+) leu1::p41nmt1-pil1-mCherry-nbr1-C(leu+)* |
| wyy814 | h+ | *his3-D1 ars1::p41nmt1-pil1-mCherry(ura+) leu1::P41nmt1-ape2-mECitrine(leu+)* |
| xm767 | | *his3-D1 leu1-32::Pnbr1-nbr1-N-TAP(leu+) ams1-13Myc::KanMx nbr1Δ::HphMX* |
| xm769 | | *his3-D1 leu1-32::Pnbr1-nbr1-C-TAP(leu+) ams1-13Myc::KanMx nbr1Δ::HphMX* |
| wyy1712 | | *his3-D1 ars1::P41nmt1-nbr1-ZZ1-L1-mCherry(ura+) leu1::P41nmt1-ams1-mECitrine(leu+)* |
| wyy1655 | h- | *his3-D1 leu1-32 ars1::P41nmt1-nbr1-ZZ1-L1-mCherry(ura+)* |
| wyy2228 | h+ | *his3-D1 ura4-D18 leu1::P41nmt1-ape4-mECitrine(leu+)* |
| wyy2272 | | *his3-D1 ars1::P41nmt1-nbr1-ZZ1-L1-mCherry(ura+) leu1::P41nmt1-ape4-mECitrine(leu+)* |
| wyy1026 | | *his3-D1 leu1-32::P41nmt1-CFP-nbr1-ZZ1Δ(leu+) ams1-mCherry::KanMx cpy1-Venus::NatMx nbr1Δ::HphMX* |
| wyy1313 | | *his3-D1 leu1-32::P41nmt1-CFP-nbr1-(ZZ2-ZZ3)Δ(leu+) ams1-mCherry::KanMx cpy1-Venus::NatMx nbr1Δ::HphMX* |
| wyy1653 | | *his3-D1 leu1-32::P41nmt1-CFP(leu+) ams1-mCherry::KanMx cpy1-Venus::NatMx nbr1Δ::HphMX* |
| wyy2417 | | *his3-D1 leu1-32::P41nmt1-CFP-nbr1-ZZ1Δ(leu+) ape4-mCherry::KanMx cpy1-Venus::NatMx nbr1Δ::NatMX ams1Δ::KanMX* |
| wyy2317 | | *his3-D1 leu1-32::P41nmt1-CFP-nbr1-(ZZ2-ZZ3)Δ(leu+) ape4-mCherry::KanMx cpy1-Venus::NatMx nbr1Δ::NatMX ams1Δ::KanMX* |
| wyy2419 | | *his3-D1 leu1-32::P41nmt1-CFP(leu+) ape4-mCherry::KanMx cpy1-Venus::NatMx nbr1Δ::NatMX ams1Δ::KanMX* |
| wyy928 | | *his3-D1 leu1-32::P41nmt1-CFP-nbr1-ZZ1Δ(leu+) lap2-mCherry::KanMx cpy1-Venus::NatMx nbr1Δ::HphMX* |
| wyy1781 | | *his3-D1 leu1-32::P41nmt1-CFP-nbr1-(ZZ2-ZZ3)Δ(leu+) lap2-mCherry::KanMx cpy1-Venus::NatMx nbr1Δ::HphMX* |
| wyy1696 | | *his3-D1 leu1-32::P41nmt1-CFP(leu+) lap2-mCherry::KanMx cpy1-Venus::NatMx nbr1Δ::HphMX* |
| wyy1316 | h- | *his3-D1 leu1-32::P41nmt1-CFP-nbr1-(ZZ2-ZZ3)Δ(leu+) ape2-mCherry::KanMx cpy1-Venus::NatMx nbr1Δ::HphMX* |
| wyy1698 | | *his3-D1 leu1-32::P41nmt1-CFP(leu+) ape2-mCherry::KanMx cpy1-Venus::NatMx nbr1Δ::HphMX* |
| wyy925 | | *his3-D1 leu1-32::P41nmt1-CFP-nbr1-ZZ1Δ(leu+) ape2-mCherry::KanMx cpy1-Venus::NatMx nbr1Δ::HphMX* |
| wyy1843 | h- | *his3-D1 ura4-D18 leu1::P1nmt1-ams1-13aa-ZZ1-ZZ2-MBP-2x3C-GFP(leu+) ams1Δ::HygMX* |
| wyy2199 | | *his3-D1 leu1::P1nmt1-ape4-13aa-ZZ1-L1-13aa-MBP-2x3C-GFP(leu+) nbr1Δ::KanMx ape4Δ::HygMX* |
| wyy2101 | h+ | *his3-D1 leu1::P41nmt1-ams1-mECitrine(leu+) ars1::P41nmt1-pil1-mCherry-ZZ1-ZZ2-D81R(ura+)* |
| wyy2288 | h+ | *his3-D1 leu1::P41nmt1-ams1-mECitrine(leu+) ars1::P41nmt1-pil1-mCherry-ZZ1-ZZ2-N63R(ura+)* |
| wyy2253 | | *his3-D1 leu1-32::P41nmt1-CFP-nbr1-(ZZ2-ZZ3)Δ-N63R(leu+) ams1-mCherry::KanMx cpy1-Venus::NatMx nbr1Δ::HphMX* |
| wyy2256 | | *his3-D1 leu1-32::P41nmt1-CFP-nbr1-(ZZ2-ZZ3)Δ-D81R(leu+) ams1-mCherry::KanMx cpy1-Venus::NatMx nbr1Δ::HphMX* |

| | | |
|---|---|---|
| wyy2138 | h- | *his3-D1 ape4-CFP::KanMx leu1::p41nmt1-pil1-mCherry-nbr1-ZZ1-ZZ2-N63R(leu+)* |
| wyy2294 | | *his3-D1 ape4-CFP::KanMx leu1::p41nmt1-pil1-mCherry-nbr1-ZZ1-ZZ2-D81R(leu+)* |
| wyy2318 | | *his3-D1 leu1-32::P41nmt1-CFP-nbr1-(ZZ2-ZZ3)Δ-N63R(leu+) ape4-mCherry::KanMx cpy1-Venus::NatMx nbr1Δ::NatMX ams1Δ::KanMX* |
| wyy2320 | | *his3-D1 leu1-32::P41nmt1-CFP-nbr1-(ZZ2-ZZ3)Δ-D81R(leu+) ape4-mCherry::KanMx cpy1-Venus::NatMx nbr1Δ::NatMX ams1Δ::KanMX* |
| wyy2545 | | *his3-D1 ars1::p41nmt1-pil1-mCherry-nbr1-ZZ1-ZZ2-P97R(ura+) leu1::P41nmt1-ams1-mECitrine(leu+)* |
| wyy2603 | | *his3-D1 leu1-32::P41nmt1-CFP-nbr1-(ZZ2-ZZ3)Δ-P97R(leu+) ams1-mCherry::KanMx cpy1-Venus::NatMx nbr1Δ::HphMX* |
| wyy2538 | h- | *his3-D1 ape4-CFP::KanMx leu1::p41nmt1-pil1-mCherry-nbr1-ZZ1-ZZ2-P97R(leu+)* |
| wyy2611 | | *his3-D1 leu1-32::P41nmt1-CFP-nbr1-(ZZ2-ZZ3)Δ-P97R(leu+) ape4-mCherry::KanMx cpy1-Venus::NatMx nbr1Δ::NatMX ams1Δ::KanMX* |
| wyy2133 | h- | *his3-D1 ape4-CFP::KanMx leu1::p41nmt1-pil1-mCherry-nbr1-ZZ1-ZZ2-L66R(leu+)* |
| wyy2298 | h- | *his3-D1 ape4-CFP::KanMx leu1::p41nmt1-pil1-mCherry-nbr1-ZZ1-ZZ2-A61R(leu+)* |
| wyy2609 | | *his3-D1 leu1-32::P41nmt1-CFP-nbr1-(ZZ2-ZZ3)Δ-L66R(leu+) ape4-mCherry::KanMx cpy1-Venus::NatMx nbr1Δ::NatMX ams1Δ::KanMX* |
| wyy2607 | | *his3-D1 leu1-32::P41nmt1-CFP-nbr1-(ZZ2-ZZ3)Δ-A61R(leu+) ape4-mCherry::KanMx cpy1-Venus::NatMx nbr1Δ::NatMX ams1Δ::KanMX* |
| wyy2544 | | *his3-D1 leu1::P41nmt1-ams1-mECitrine(leu+) ars1::P41nmt1-pil1-mCherry-ZZ1-ZZ2-L66R(ura+)* |
| wyy2543 | | *his3-D1 leu1::P41nmt1-ams1-mECitrine(leu+) ars1::P41nmt1-pil1-mCherry-ZZ1-ZZ2-A61R(ura+)* |
| wyy2602 | | *his3-D1 leu1-32::P41nmt1-CFP-nbr1-(ZZ2-ZZ3)Δ-L66R(leu+) ams1-mCherry::KanMx cpy1-Venus::NatMx nbr1Δ::HphMX* |
| wyy2600 | | *his3-D1 leu1-32::P41nmt1-CFP-nbr1-(ZZ2-ZZ3)Δ-A61R(leu+) ams1-mCherry::KanMx cpy1-Venus::NatMx nbr1Δ::HphMX* |
| wyy2103 | h+ | *his3-D1 leu1::P41nmt1-ams1-T2A-mECitrine(leu+) ars1::P41nmt1-pil1-mCherry-ZZ1-ZZ2(ura+)* |
| wyy2566 | h+ | *his3-D1 leu1::P41nmt1-ams1-L3A-mECitrine(leu+) ars1::P41nmt1-pil1-mCherry-ZZ1-ZZ2(ura+)* |
| wyy2569 | h+ | *his3-D1 leu1::P41nmt1-ams1-F4A-mECitrine(leu+) ars1::P41nmt1-pil1-mCherry-ZZ1-ZZ2(ura+)* |
| wyy2405 | h+ | *his3-D1 leu1::P41nmt1-pil1-mCherry-ZZ1-ZZ2(leu+) ars1::P41nmt1-pmt3(1-109)-ape4-mECitrine(ura+)* |
| wyy2407 | h+ | *his3-D1 leu1::P41nmt1-pil1-mCherry-ZZ1-ZZ2(leu+) ars1::P41nmt1-pmt3(1-111)-ape4-mECitrine(ura+)* |
| wyy2408 | h+ | *his3-D1 leu1::P41nmt1-pil1-mCherry-ZZ1-ZZ2(leu+) ars1::P41nmt1-pmt3(1-111)-ape4-M1A-mECitrine(ura+)* |
| wyy2410 | h+ | *his3-D1 leu1::P41nmt1-pil1-mCherry-ZZ1-ZZ2(leu+) ars1::P41nmt1-pmt3(1-111)-ape4-Q2A-mECitrine(ura+)* |
| wyy2394 | h+ | *his3-D1 ura4-D18 leu1::P41nmt1-pmt3(1-109)-ape4-mECitrine(leu+)* |
| wyy2397 | h+ | *his3-D1 ura4-D18 leu1::P41nmt1-pmt3(1-111)-ape4-mECitrine(leu+)* |
| wyy2400 | h+ | *his3-D1 ura4-D18 leu1::P41nmt1-pmt3(1-111)-ape4-M1A-mECitrine(leu+)* |
| wyy2403 | h+ | *his3-D1 ura4-D18 leu1::P41nmt1-pmt3(1-111)-ape4-Q2A-mECitrine(leu+)* |

| | | |
|---|---|---|
| wyy2312 | h+ | *his3-D1 leu1::P41nmt1-pil1-mCherry-ZZ1-ZZ2(leu+) ars1::P41nmt1-ape4-mECitrine(ura+)* |
| wyy2444 | h+ | *his3-D1 leu1::P41nmt1-pil1-mCherry-ZZ1-ZZ2(leu+) ars1::P41nmt1-ape4-L3A-mECitrine(ura+)* |
| wyy2699 | h+ | *his3-D1 leu1::P41nmt1-pil1-mCherry-ZZ1-ZZ2(leu+) ars1::P41nmt1-ape4-M7R-mECitrine(ura+)* |
| wyy2702 | h+ | *his3-D1 leu1::P41nmt1-pil1-mCherry-ZZ1-ZZ2(leu+) ars1::P41nmt1-ape4-A9R-mECitrine(ura+)* |
| wyy2705 | h+ | *his3-D1 leu1::P41nmt1-pil1-mCherry-ZZ1-ZZ2(leu+) ars1::P41nmt1-ape4-K12E-mECitrine(ura+)* |
| wyy2593 | h+ | *his3-D1 leu1::P41nmt1-pil1-mCherry-ZZ1-ZZ2(leu+) ars1::P41nmt1-pmt3(1-111)-ams1(1-1077)-mECitrine(ura+)* |
| wyy2743 | h+ | *his3-D1 leu1::P41nmt1-pil1-mCherry-ZZ1-ZZ2(leu+) ars1::P41nmt1-pmt3(1-111)-ams1(2-1077)-mECitrine(ura+)* |
| wyy949 | h+ | *his3-D1 ura4-D18 leu1::Pnmt1-ams1-FFH(leu+)* |
| wyy2314 | h+ | *his3-D1 leu1::Pnmt1-ape4-FFH(leu+) ape4Δ::KanMX* |
| wyy2573 | | *his3-D1 leu1-32 ars1::P41nmt1-CFP-nbr1-(ZZ2-ZZ3)D(ura4+) ams1-mCherry::KanMx Cpy1-Venus::NatMx nbr1Δ::HphMX ape4Δ::KanMX* |
| wyy2501 | | *his3-D1 leu1::Pnmt1-ape4-FFH(leu+) ars1::P41nmt1-CFP-nbr1-(ZZ2-ZZ3)D(ura+) ams1-mCherry::KanMx Cpy1-Venus::NatMx nbr1Δ::HphMX ape4Δ::KanMX* |
| wyy2269 | h+ | *his3-D1 leu1::P41nmt1-pil1-mCherry-ZZ1-ZZ2-K100A(leu+) ars1::P41nmt1-ams1-mECitrine(ura+)* |
| wyy2290 | h+ | *his3-D1 leu1::P41nmt1-pil1-mCherry-ZZ1-ZZ2-F80A(leu+) ars1::P41nmt1-ams1-mECitrine(ura+)* |
| wyy2136 | h- | *his3-D1 leu1::P41nmt1-pil1-mCherry-ZZ1-ZZ2-F80A(leu+) ape4-CFP::KanMX* |
| wyy2131 | h- | *his3-D1 leu1::P41nmt1-pil1-mCherry-ZZ1-ZZ2-K100A(leu+) ape4-CFP::KanMX* |

Table S3. Plasmids used in this study.

| Plasmid | Descriptive name | Description |
|---|---|---|
| pDB3389 | pDUAL-Pnmt1-Nbr1-YFH | Matsuyama et al. 2006 |
| pDB3403 | pJK148-Pnbr1-Nbr1-TAP | Liu et al. 2015 |
| pWYY56 | pDUAL-P41nmt1-Ams1-mECitrine | pDUAL plasmid expressing Ams1-mECitrine from P41nmt1 promoter |
| pWYY534 | pDUAL-P41nmt1-GBP-3xUb | pDUAL plasmid expressing GBP-3xUb from P41nmt1 promoter |
| pWYY183 | pDUAL-P41nmt1-mCherry-GBP-3xUb | pDUAL plasmid expressing mCherry-GBP-3xUb from P41nmt1 promoter |
| pWYY83 | pDUAL-P41nmt1-Pil1-mCherry-Nbr1-N | pDUAL plasmid expressing Pil1-mCherry-Nbr1-N from P41nmt1 promoter |
| pWYY535 | pDUAL-P41nmt1-Pil1-mCherry-Nbr1-ZZ1 | pDUAL plasmid expressing Pil1-mCherry-Nbr1-ZZ1 from P41nmt1 promoter |
| pWYY119 | pDUAL-P41nmt1-Pil1-mCherry-Nbr1-ZZ1-ZZ2 | pDUAL plasmid expressing Pil1-mCherry-Nbr1-ZZ1-ZZ2 from P41nmt1 promoter |
| pWYY199 | pDUAL-P41nmt1-Pil1-mCherry-Nbr1-ZZ2-ZZ3 | pDUAL plasmid expressing Pil1-mCherry-Nbr1-ZZ2-ZZ3 from P41nmt1 promoter |
| pWYY197 | pDUAL-P41nmt1-Pil1-mCherry-Nbr1-ZZ1-L1 | pDUAL plasmid expressing Pil1-mCherry-Nbr1-ZZ1-L1 from P41nmt1 promoter |
| pWYY159 | pDUAL-P41nmt1-Pil1-mCherry-Nbr1-L1-ZZ2 | pDUAL plasmid expressing Pil1-mCherry-Nbr1-L1-ZZ2 from P41nmt1 promoter |
| pWYY166 | pDUAL-P41nmt1-Pil1-mCherry-Nbr1-L2-ZZ3 | pDUAL plasmid expressing Pil1-mCherry-Nbr1-L2-ZZ3 from P41nmt1 promoter |
| pWYY100 | pDUAL-P41nmt1-Pil1-mCherry-Nbr1-C | pDUAL plasmid expressing Pil1-mCherry-Nbr1-C from P41nmt1 promoter |
| pWYY59 | pDUAL-P41nmt1-Pil1-mCherry | pDUAL plasmid expressing Pil1-mCherry from P41nmt1 promoter |
| pWYY114 | pDUAL-P41nmt1-Lap2-mECitrine | pDUAL plasmid expressing Lap2-mECitrine from P41nmt1 promoter |
| pWYY117 | pDUAL-P41nmt1-Ape2-mECitrine | pDUAL plasmid expressing Ape2-mECitrine from P41nmt1 promoter |
| pWYY234 | pDUAL-P41nmt1-Nbr1-ZZ1-L1-mCherry | pDUAL plasmid expressing Nbr1-ZZ1-L1-mCherry from P41nmt1 promoter |
| pWYY400 | pDUAL-P41nmt1-Ape4-mECitrine | pDUAL plasmid expressing Ape4-mECitrine from P41nmt1 promoter |
| pXM26 | pDUAL-P41nmt1-CFP-Nbr1-ΔZZ1 | pDUAL plasmid expressing CFP-Nbr1-ΔZZ1 from P41nmt1 promoter |
| pWYY209 | pDUAL-P41nmt1-CFP-Nbr1-Δ(ZZ2-ZZ3) | pDUAL plasmid expressing CFP-Nbr1-Δ(ZZ2-ZZ3) from P41nmt1 promoter |
| pWYY208 | pDUAL-P41nmt1-CFP | pDUAL plasmid expressing CFP from P41nmt1 promoter |
| pWYY295 | pDUAL-Pnmt1-Ams1-13aa-Nbr1-ZZ1-ZZ2-MBP-2x3C-GFP | pDUAL plasmid expressing Ams1-13aa-Nbr1-ZZ1-ZZ2-MBP-2x3C-GFP from Pnmt1 promoter |

| | | |
|---|---|---|
| pWYY406 | pDUAL-Pnmt1-Ape4-13aa-Nbr1-ZZ1-L1-13aa-MBP-2x3C-GFP | pDUAL plasmid expressing Ape4-13aa-Nbr1-ZZ1-L1-13aa-MBP-2x3C-GFP from Pnmt1 promoter |
| pWYY337 | pDUAL-P41nmt1-Pil1-mCherry-Nbr1-ZZ1-ZZ2-D81R | pDUAL plasmid expressing Pil1-mCherry-Nbr1-ZZ1-ZZ2-D81R from P41nmt1 promoter |
| pWYY342 | pDUAL-P41nmt1-Pil1-mCherry-Nbr1-ZZ1-ZZ2-N63R | pDUAL plasmid expressing Pil1-mCherry-Nbr1-ZZ1-ZZ2-N63R from P41nmt1 promoter |
| pWYY420 | pDUAL-P41nmt1-CFP-Nbr1-Δ(ZZ2-ZZ3)-N63R | pDUAL plasmid expressing CFP-Nbr1-Δ(ZZ2-ZZ3)-N63R from P41nmt1 promoter |
| pWYY419 | pDUAL-P41nmt1-CFP-Nbr1-Δ(ZZ2-ZZ3)-D81R | pDUAL plasmid expressing CFP-Nbr1-Δ(ZZ2-ZZ3)-D81R from P41nmt1 promoter |
| pWYY531 | pDUAL-P41nmt1-Pil1-mCherry-Nbr1-ZZ1-ZZ2-P97R | pDUAL plasmid expressing Pil1-mCherry-Nbr1-ZZ1-ZZ2-P97R from P41nmt1 promoter |
| pWYY513 | pDUAL-P41nmt1-CFP-Nbr1-Δ(ZZ2-ZZ3)-P97R | pDUAL plasmid expressing CFP-Nbr1-Δ(ZZ2-ZZ3)-P97R from P41nmt1 promoter |
| pWYY345 | pDUAL-P41nmt1-Pil1-mCherry-Nbr1-ZZ1-ZZ2-L66R | pDUAL plasmid expressing Pil1-mCherry-Nbr1-ZZ1-ZZ2-L66R from P41nmt1 promoter |
| pWYY344 | pDUAL-P41nmt1-Pil1-mCherry-Nbr1-ZZ1-ZZ2-A61R | pDUAL plasmid expressing Pil1-mCherry-Nbr1-ZZ1-ZZ2-A61R from P41nmt1 promoter |
| pWYY512 | pDUAL-P41nmt1-CFP-Nbr1-Δ(ZZ2-ZZ3)-L66R | pDUAL plasmid expressing CFP-Nbr1-Δ(ZZ2-ZZ3)-L66R from P41nmt1 promoter |
| pWYY509 | pDUAL-P41nmt1-CFP-Nbr1-Δ(ZZ2-ZZ3)-A61R | pDUAL plasmid expressing CFP-Nbr1-Δ(ZZ2-ZZ3)-A61R from P41nmt1 promoter |
| pWYY351 | pDUAL-P41nmt1-Ams1-T2A-mECitrine | pDUAL plasmid expressing Ams1-T2A-mECitrine from P41nmt1 promoter |
| pWYY352 | pDUAL-P41nmt1-Ams1-L3A-mECitrine | pDUAL plasmid expressing Ams1-L3A-mECitrine from P41nmt1 promoter |
| pWYY463 | pDUAL-P41nmt1-Ams1-F4A-mECitrine | pDUAL plasmid expressing Ams1-F4A-mECitrine from P41nmt1 promoter |
| pWYY437 | pDUAL-P41nmt1-Pmt3(1-109)-Ape4-mECitrine | pDUAL plasmid expressing Pmt3(1-109)-Ape4-mECitrine from P41nmt1 promoter |
| pWYY439 | pDUAL-P41nmt1-Pmt3(1-111)-Ape4-mECitrine | pDUAL plasmid expressing Pmt3(1-111)-Ape4-mECitrine from P41nmt1 promoter |
| pwyy442 | pDUAL-P41nmt1-Pmt3(1-111)-Ape4-M1A-mECitrine | pDUAL plasmid expressing Pmt3(1-111)-Ape4-M1A-mECitrine from P41nmt1 promoter |
| pWYY443 | pDUAL-P41nmt1-Pmt3(1-111)-Ape4-Q2A-mECitrine | pDUAL plasmid expressing Pmt3(1-111)-Ape4-Q2A-mECitrine from P41nmt1 promoter |
| pWYY475 | pDUAL-P41nmt1-Ape4-L3A-mECitrine | pDUAL plasmid expressing Ape4-L3A-mECitrine from P41nmt1 promoter |
| pWYY559 | pDUAL-P41nmt1-Ape4-M7R-mECitrine | pDUAL plasmid expressing Ape4-M7R-mECitrine from P41nmt1 promoter |
| pWYY561 | pDUAL-P41nmt1-Ape4-A9R-mECitrine | pDUAL plasmid expressing Ape4-A9R-mECitrine from P41nmt1 promoter |
| pWYY563 | pDUAL-P41nmt1-Ape4-K12E-mECitrine | pDUAL plasmid expressing Ape4-K12E-mECitrine from P41nmt1 promoter |
| pWYY508 | pDUAL-P41nmt1-Pmt3(1-111)-Ams1(1-1077)-mECitrine | pDUAL plasmid expressing Pmt3(1-111)-Ams1(1-1077)-mECitrine from P41nmt1 promoter |
| pWYY555 | pDUAL-P41nmt1-Pmt3(1-111)-Ams1(2-1077)-mECitrine | pDUAL plasmid expressing Pmt3(1-111)-Ams1(2-1077)-mECitrine from P41nmt1 promoter |

| | | |
|---|---|---|
| pWYY135 | pDUAL-Pnmt1-Ams1-FFH | pDUAL plasmid expressing Ams1-FFH from Pnmt1 promoter |
| pWYY399 | pDUAL-Pnmt1-Ape4-FFH | pDUAL plasmid expressing Ape4-FFH from Pnmt1 promoter |
| pWYY363 | pETDuet-His6-GST-Nbr1-ZZ1-L1 | pETDuet plasmid expressing His6-GST-Nbr1-ZZ1-L1 from T7/lac promoter |
| pWYY311 | pDUAL-P41nmt1-Pil1-mCherry-Nbr1-ZZ1-ZZ2-F80A | pDUAL plasmid expressing Pil1-mCherry-Nbr1-ZZ1-ZZ2-F80A from P41nmt1 promoter |
| pWYY309 | pDUAL-P41nmt1-Pil1-mCherry-Nbr1-ZZ1-ZZ2-K100A | pDUAL plasmid expressing Pil1-mCherry-Nbr1-ZZ1-ZZ2-K100A from P41nmt1 promoter |
